# SOX11 stimulates γδ T-cell differentiation and synergizes with LMO2 and MYCN to drive γδ-like T-cell acute lymphoblastic leukemia

**DOI:** 10.64898/2026.07.28.741231

**Authors:** André Almeida, Igor Fijalkowski, Kasper Thorhauge Christensen, Herlinde De Waele, Petri Pölönen, Marlies Vanden Bempt, Els Van Ammel, Sara T’Sas, Béatrice Lintermans, Panagiotis Ntziachristos, Kaat Durinck, Frank Speleman, Tom Taghon, Jan Cools, Charles G. Mullighan, David Teachey, Pieter Van Vlierberghe, Tim Pieters, Steven Goossens

## Abstract

T-cell acute lymphoblastic leukemia (T-ALL) is a heterogeneous hematologic malignancy in which LMO2 γδ-like T-ALL represents a rare but clinically aggressive subtype associated with poor treatment response and inferior survival. Integrated transcriptomic analyses identified high SOX11 expression as a defining feature of high-risk LMO2 γδ-like T-ALL, where elevated SOX11 levels correlated with refractory disease and poor clinical outcome. To investigate the functional role of SOX11 in γδ T-cell biology and leukemogenesis, we generated a conditional *R26-SOX11* mouse model enabling lineage-specific SOX11 overexpression in T-cell progenitors. SOX11 expression promoted expansion of the innate γδ T-cell compartment in thymus, spleen, and bone marrow, accompanied by transcriptional activation of γδ T-cell differentiation, activation, and cytotoxicity programs. However, SOX11 overexpression alone was insufficient to induce leukemia or confer thymocyte self-renewal capacity. In contrast, combined SOX11 and LMO2 overexpression markedly accelerated T-ALL development and strongly increased the incidence of γδ-like leukemias, thereby recapitulating the human high-risk LMO2 γδ-like T-ALL subtype. Mechanistically, SOX11 expanded the pre-leukemic DN3 thymocyte compartment in LMO2-driven mouse model while promoting differentiation toward the γδ lineage. Transcriptomic profiling identified activation of MYCN-associated transcriptional programs in SOX11/LMO2 pre-leukemic thymocytes. Consistently, MYCN was highly expressed in human LMO2 γδ-like T-ALL, and recurrent stabilizing MYCN P44L mutations were enriched in this subtype. Functional validation using genetic and transplantation-based mouse models demonstrated that SOX11 cooperates with MYCN to accelerate T-ALL onset. Together, these findings establish a cooperative SOX11–MYCN oncogenic axis driving γδ-like T-ALL and provide a novel preclinical model for investigating therapeutic vulnerabilities in this high-risk leukemia subtype.

## Introduction

T-cell acute lymphoblastic leukemia (T-ALL) is an aggressive hematological malignancy characterized by a diffuse infiltration of T-lymphoblasts in the bone marrow ^1^. T-ALL accounts for approximately 15% of pediatric and 25% of adult acute lymphoblastic leukemia (ALL) cases ^2,3^. Although the use of intensified chemotherapy and bone marrow transplantation increased survival rates, still 10-15% of pediatric and 40-45% of adult T-ALL patients exhibit primary resistance to standard therapy, or eventually relapse ^4,5^. Effective salvage treatment options are lacking for these patients, resulting in dismal outcomes. Identifying patients at high risk for resistance/recurrence and developing novel therapeutic strategies are critical priorities.

T-ALL is a highly heterogenous disease. Approximately 10% of T-ALL cases originate from the malignant transformation of thymic precursors committed to the γδ T-cell lineage ^6–9^, a minor subset of thymic T cells defined by expression of a γδ T-cell receptor (TCR). Unlike the conventional αβ T-cells, γδ T-cells recognize antigens in a largely MHC-independent manner, allowing them to bridge innate and adaptive immunity. Although γδ T-ALL shares many features with conventional αβ T-ALL, accumulating evidence indicates that it represents a biologically distinct and heterogeneous group. Although γδ T-ALL was initially associated with an unfavorable clinical outcome, with particularly poor prognosis reported in young children (<3 years of age) ^7,8^, these findings could not be confirmed in an independent cohort ^6,9^. This discrepancy highlights the need for a more refined biological classification of γδ T-ALLs. Comprehensive molecular profiling of 200 γδ T-ALL cases revealed that clinical outcome is strongly influenced by the underlying genetic drivers ^7^. High-risk cases are enriched for genetic alterations resulting in LMO2 activation and STAG2 inactivation. In addition, γδ T-ALL can be stratified according to the activation of homeodomain (HD)-containing transcription factor, including members of the HOXA cluster and NK-like homeobox family. These cortical-like HD^+^ γδ T-ALLs exhibit a more favorable initial response to treatment compared to *bona fide* HD^−^ γδ T-ALL, although no statistical difference in long-term outcome was observed ^9^. An integrated multivariate analysis identified 15 molecular subtypes of pediatric T-ALL distinguished by their genomic drivers, gene expression, developmental states, and clinical outcomes ^10^. Among these, the LMO2 γδ-like subtype, which is reminiscent of *bona fide* HD^−^ γδ T-ALL, was associated with induction failure and poor prognosis. Together, these findings highlight an urgent need to develop more effective treatment strategies for high-risk γδ T-ALL. However, the molecular mechanisms underlying the aggressive clinical course of this rare subtype of patients remains incompletely understood.

*SOX11* is a single-exon gene that encodes a pioneering transcriptional activator ^11–13^. During mouse embryogenesis, *Sox11* is prominently expressed in neural progenitors together with the other SOX-C family members, *Sox4* and *Sox12* ^11^. High *Sox11* expression is also detected in developing bone, the digestive tract, genital and endocardial tissues ^11^. In these embryonic tissues, SOX11 plays key roles in progenitor cell differentiation, proliferation and survival. Consistently, *Sox11*-null mice exhibit severe developmental defects and die shortly after birth ^11^. Although *Sox11* expression is largely absent from adult mouse tissues, aberrant *SOX11* expression has been reported in several human cancers, including neuroblastoma and other neural-derived tumors, breast cancer, epithelial ovarian cancer and gastrointestinal malignancies ^11,14–19^. Depending on the tumor context, SOX11 expression has been associated with either oncogenic or tumor-suppressor functions, influencing proliferation, metastatic potential and patient prognosis ^11,17^. In hematologic malignancies, SOX11 is a well-established diagnostic and prognostic biomarker in mantle cell lymphoma (MCL) ^20,21^, and aberrant SOX11 expression has also been detected in B-cell precursor ALL ^22^. Although SOX11 is absent from normal T-cells, ^23–25^, its overexpression has been consistently observed in a subset of T-cell malignancies ^21,22^. However, its functional role, prognostic relevance, and potential as a therapeutic target in T-cell leukemogenesis remain poorly understood.

Here, we identify SOX11 as a key molecular determinant of high-risk LMO2 γδ-like T-ALL. Through the generation of a conditional transgenic mouse model and CITEseq analysis, we uncover a previously unrecognized role for SOX11 in promoting innate γδ T-cell expansion and demonstrate its potent cooperation with LMO2 in driving γδ-like leukemogenesis. Our findings further reveal MYCN as a critical downstream effector of this program, defining a SOX11– MYCN oncogenic axis that contributes to disease initiation and progression. Collectively, this work establishes the first faithful mouse model of LMO2 γδ-like T-ALL and provides a mechanistic framework for understanding the aggressive clinical behavior of this leukemia subtype, opening new avenues for biomarker development and targeted therapeutic intervention.

## Results

### *SOX11* is highly expressed in LMO2 γδ-like T-ALL

γδ T-ALL represents approximately 10% of T-ALL cases and is associated with a higher incidence of refractory disease and an unfavorable prognosis, particularly in very young children (<3 years) ^7^. While γδ T-cells account for ∼1% of thymocytes, γδ T-ALL represent ∼10% of T-ALL cases ^7,26^, suggesting the presence of an amplifying oncogenic mechanism. Recent classification of T-ALLs using both genomic and transcriptomic features, identified LMO2 γδ-like T-ALL as a high-risk subtype (Fig. S1A) ^10^. To identify oncogenic transcription factors (TFs) potentially involved in this mechanism, we assessed which ones were overrepresented in the LMO2 γδ-like subtype compared to all other T-ALL subtypes. This analysis identified SOX13 and SOX11 as the top candidates (Fig. 1A,B; Fig. S1B). While the role of SOX13 in γδ T-cell development has been documented ^27^, the contribution of SOX11 to γδ T-cell biology and γδ T-ALL development remains unknown. Of note, SOX11 is also expressed in select other T-ALL subtypes, including STAG-LMO2, KMT2A, and TAL1 αβ-like leukemias; however, its expression does not correlate with γδ TCR status or induction failure outside the LMO2 γδ-like subtype (Fig. 1C,D; Fig. S1C,D). Moreover, within the LMO2 γδ-like subtype, high SOX11 expression is associated with inferior clinical outcome (Fig. 1E). In contrast, SOX11 expression was not prognostic in other T-ALL subtypes, with the exception of TAL1 αβ-like T-ALL, in which elevated SOX11 expression correlated with a favorable outcome (Fig. 1SE). Overall, SOX11 represents a prognostic marker in LMO2 γδ-like T-ALL, with elevated expression correlating with induction failure and poor prognosis, thereby defining this aggressive leukemia subtype.

**Figure 1.**
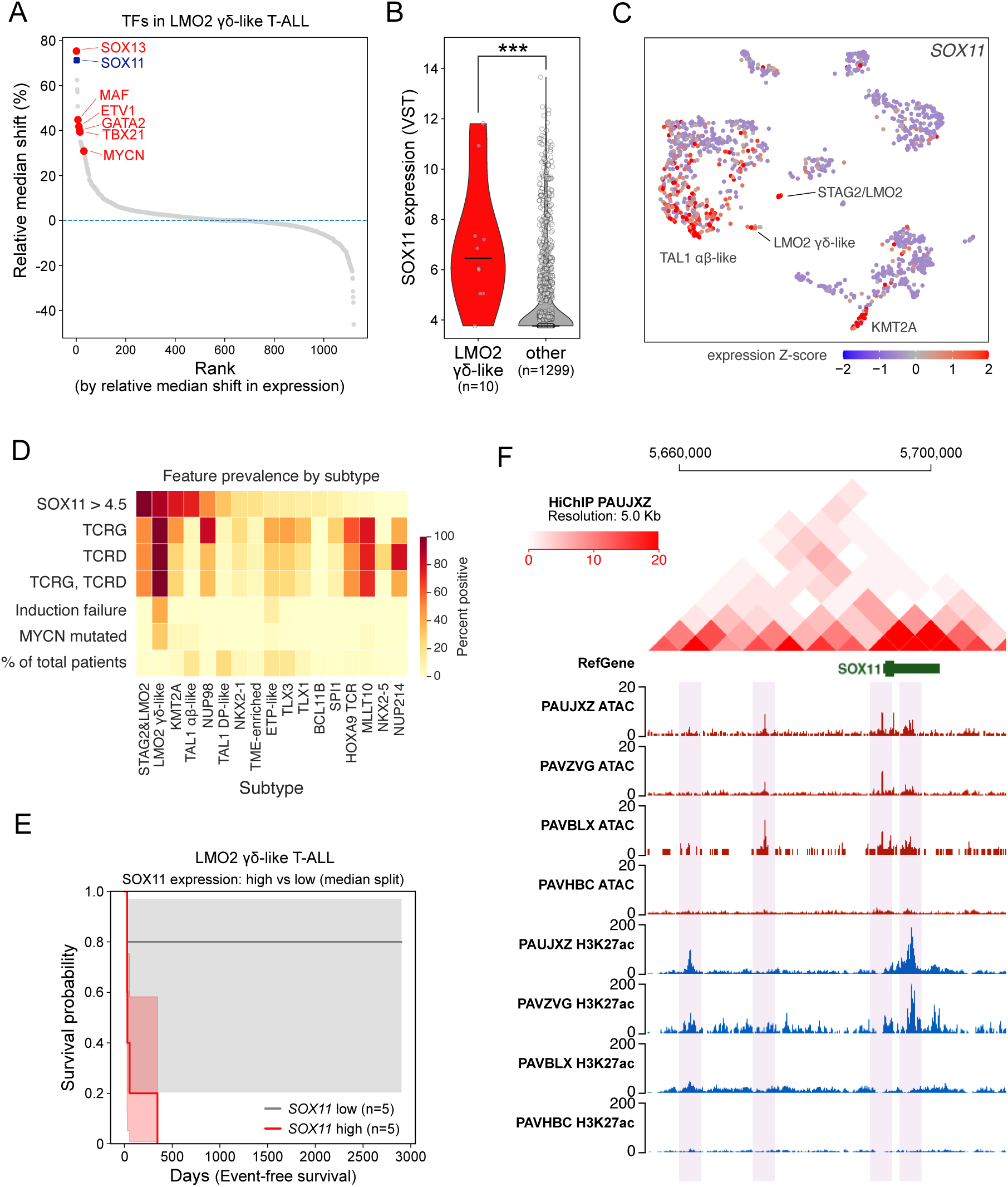
SOX11 marks aggressive LMO2 γδ-like T-ALL associated with induction failure and poor outcome. **(A)** Waterfall plot showing the relative median expression shift of transcription factors in LMO2 γδ-like T-ALL compared with all other T-ALL subtypes, based on variance-stabilizing transformed (VST) RNA-seq data from 1309 patients ^10^. Median expression was calculated for each gene in LMO2 γδ-like cases and compared to the median level of expression for all other subtypes. Selected transcription factors are highlighted in red and SOX11 in blue. **(B)** Violin plot depicting *SOX11* expression in LMO2 γδ-like T-ALL (n=10) compared with all other T-ALL subtypes (n=1293) ^10^. Significance was assessed using a two-sided Mann-Whitney U test (p = 1.74 × 10^−4^). **(C)** UMAP scatterplot of T-ALL subtypes showing SOX11 expression. **(D)** Heatmap showing the prevalence of selected clinical, molecular, and immunogenetic features across T-ALL subtypes^10^. Columns represent molecular subtypes, ordered by decreasing prevalence of SOX11-high cases. Rows indicate the percentage of cases within each subtype with SOX11 expression > 4.5 VST (threshold for SOX11 expression at all), TCRG rearrangement, TCRD rearrangement, combined TCRG/TCRD rearrangement, induction failure, or MYCN mutation. The bottom row shows the proportion of the total cohort assigned to each subtype. SOX11 positivity was defined as VST expression > 4.5. This cutoff was selected based on the density distribution of SOX11 expression. TCRG and TCRD indicate cases with any rearrangement involving the respective locus. TCRG, TCRD indicates cases with both gamma and delta rearrangements. Color intensity represents percentage positive within each subtype, except for “% of total patients,” which indicates subtype frequency in the full cohort. **(E)** Kaplan-Meier analysis of event-free survival in LMO2 γδ-like T-ALL cases ^10^ stratified by SOX11 expression using a within-subtype median split. Groups were compared by log-rank test (*, *P* = 0.021). **(F)** Genome browser view of the *SOX11* locus showing H3K27ac HiChIP interactions in the SOX11-expressing T-ALL patient sample PAUJXZ (top panel), together with ATAC-seq and H3K27ac ChIP-seq profiles from SOX11^hi^ and SOX11^low^/negative T-ALL patient samples (bottom panels). PAVBLX sample is LMO2 γδ-like sample, while PAVZVG is STAG2/LMO2 and PAUJXZ and PAVHBC are TAL1 DP-like.

To investigate the regulatory mechanisms associated with *SOX11* expression in T-ALL, we analyzed H3K27ac HiChIP and ATAC-seq chromatin accessibility profiles from 20 T-ALL patient samples^10,28^. This analysis identified four candidate *SOX11*-associated regulatory elements with evidence of enhancer activity in T-ALL, including in an LMO2 γδ-like case with elevated *SOX11* expression (Fig. 1F, Fig. S1F). Collectively, these findings nominate a set of putative cis-regulatory elements that may contribute to aberrant SOX11 expression in T-ALL.

### SOX11 overexpression stimulates γδ T-cell differentiation

To investigate the role of SOX11 in γδ T-cell development and the oncogenic transformation to γδ-T-ALL, we generated a conditional SOX11 overexpression mouse model (*R26-SOX11^tg/tg^* mice; Fig. 2A). To validate this novel transgenic line, these mice were crossed with *CD2-iCre^tg^* mice, which express Cre recombinase in early lymphoid progenitors. This yielded *R26-SOX11^tg/tg^; CD2-iCre^tg/+^* mice (hereafter referred to as *SOX11^CD2^*) and Cre negative *R26-SOX11^tg/tg^; CD2-iCre^+/+^* littermates (referred to as *controls*). Upon Cre-mediated excision of the floxed transcriptional stop cassette, consistent conditional expression of a bicistronic transgenic transcript was detected in bone marrow of *SOX11^CD2^*, encoding both SOX11 and the eGFP reporter (Fig. S2A,B). Flow cytometric analysis revealed that SOX11 overexpression increased the percentages of CD4^−^CD8^−^CD3^+^TCRγδ^+^ γδ T-cells in the thymus, spleen and bone marrow of *SOX11^CD2^* mice (Fig. 2B,C). The increase of γδ T-cells was confirmed by CITE-seq on Cre-negative controls (n=3) and *SOX11^CD2^* mice (n=3) (Fig. 2D,E and Fig. S2C-F).

**Figure 2.**
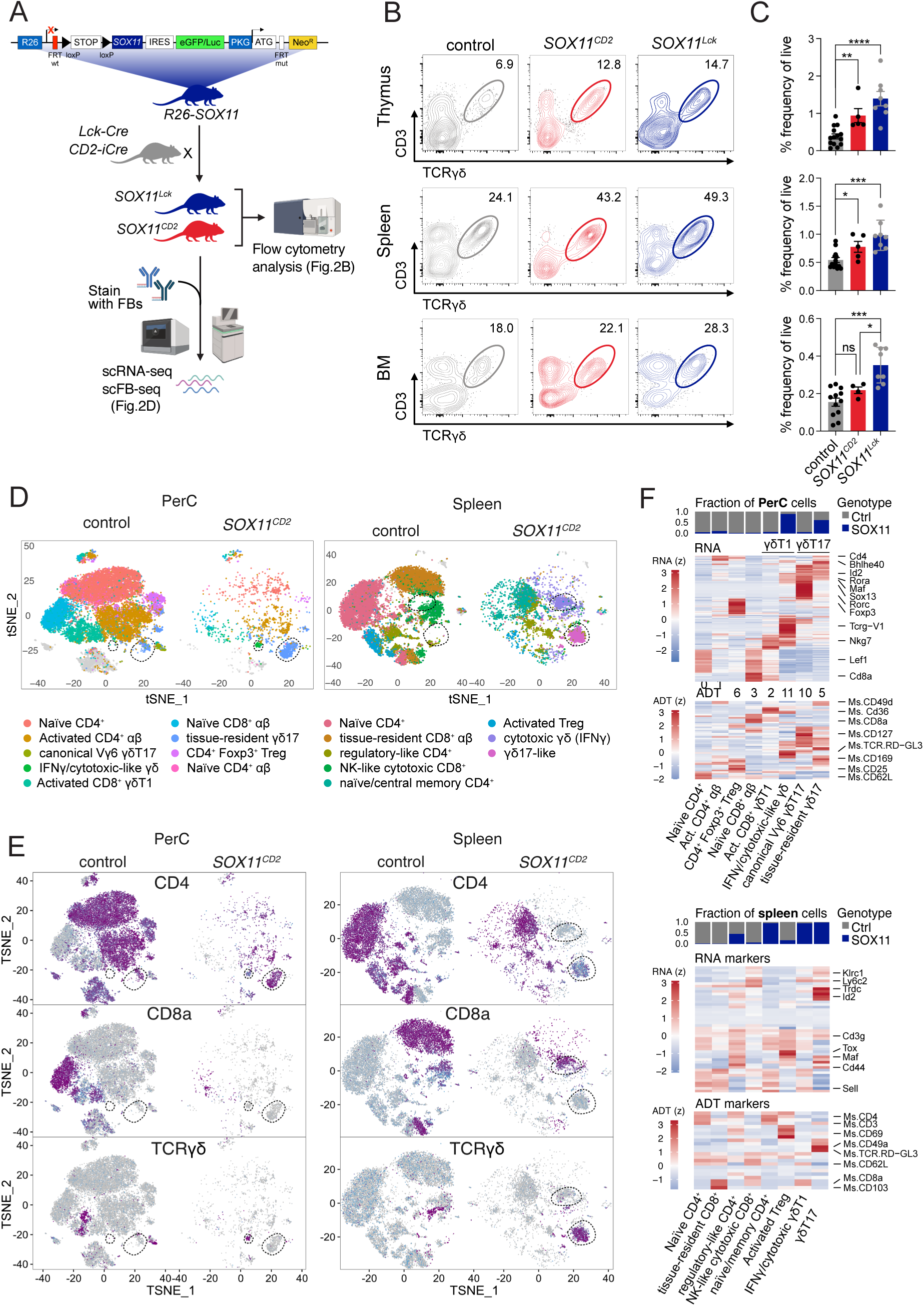
SOX11 blocks adaptive immunity and promotes γδ T-cell formation. **(A)** Schematic overview of the *R26-SOX11* mouse model enabling conditional SOX11 overexpression, the breeding strategies used to induce SOX11 expression in lymphoid progenitors (via *CD2-iCre*) or T-cell progenitors (via *Lck-Cre*), and the experimental workflow. The study design included flow cytometric analysis of both Cre-driven models, as well as single cell RNA sequencing (scRNA-seq) and single cell feature barcodes sequencing (scFB-seq) performed on cells isolated from three *R26-SOX11^tg/tg^;CD2-iCre^tg/+^ (SOX11^CD2^)* mice and three Cre-negative littermate controls. FRT, flippase recognition target; R26: ROSA26; IRES: independent ribosomal entry site; PKG, phosphoglycerate kinase 1; NeoR, neomycin resistance gene; Luc, luciferase. **(B)** Flow cytometric analysis of thymocytes, splenocytes or bone marrow (BM) cells from 8-week-old Cre-negative littermate controls or from mice expressing SOX11 in common lymphoid progenitors (*SOX11^CD2^*) or thymic progentoris *(SOX11^Lck^)*. γδ T-cells were defined as CD3^+^TCRγδ ^+^ cells within the CD4^−^CD8^−^ DN thymocyte population, pregated on single, live CD90.2^+^ cells. **(C)** Bar graphs displaying frequencies of γδ T-cells (as a percentage of live cells) from flow cytometry data shown in (F). * *P* < 0.05, ** *P* <0.01, *** *P* < 0.001, **** *P* <0.0001. ns: not significant. **(D)** t-SNE plot showing annotated single-cell clusters from control and SOX11-expressing (*SOX11^CD2^)* cells from spleen or peritoneal cavity (PerC). Clusters were defined based on surface marker expression derived from FB-seq. Dotted lines indicate clusters enriched or specifically induced upon SOX11 expression. **(E)** (top) Bar graph showing, for each cluster, the relative proportions of control and SOX11-expressing (*SOX11^CD2^)* cells. Heatmaps showing differentially expressed cell surface markers (middle) and genes (middle) in different clusters. ADT: Antibody-determining tag.

In mice, two major γδ T-cell subsets have been described: IL-17-producing γδT17 cells, which preferentially reside in the peritoneal cavity (PerC), dermis, and adipose tissue, and IFNγ-producing γδT1 cells, which are enriched in the skin, liver, and spleen ^29^. To determine which subsets are induced upon SOX11 overexpression, we mined CITE-seq data on PerC, spleen and bone marrow (BM) from Cre-negative controls (n=3) and *SOX11^CD2^* mice (n=3), representing tissues enriched for γδT17 and γδT1 cells. In the PerC of control mice, we identified a canonical Vγδ6^+^ γδT17 population characterized by expression of *Tcrg-V6*, *Sox13*, *Rorc, Il23r, Maf, Cxcr6, Il17re*. In contrast, SOX11-induced γδT17 cells detected across PerC, spleen and, BM, exhibited reduced expression of key γδT17-defining genes, including *Rorc, SOX13, Il23r, and Il17re*. Instead, these cells acquired an activated and partially reprogrammed phenotype characterized by increased expression of activation-associated genes (*Tnfrsf4*, *Icos* and *Ly6a*), tissue residency factors (*Cxcr6, Rora* and *Id2*), and γδT1/NK-associated features (*Nkg7, Klrd1, Ifngr1,* Id2)(Fig. 2F, Fig. S2-3).

To further assess the extent of this transcriptional shift, we compared our data with published γδT17 and γδT1 reference datasets generated using endogenous fluorescent reporters inserted into the *Il17a* and *Ifng* loci, enabling purification and transcriptional profiling of bona fide γδT17 and γδT1 populations ^30^. Wild-type PerC γδT17 cells showed strong concordance with the reference γδ17 signature, sharing 280 genes, including canonical γδT17 markers (*Rorc*, *Il23r*, *Sox13*, *Il7r)*, and additional genes previously validated using this reporter-based system, such as *Adam12*, *Ret*, *Tmem176b*, *Cysltr1*, and *Serpinb1a* (Fig. S3C). In contrast, SOX11-induced PerC γδT17 cells upregulated multiple genes characteristic of the γδ1 program, including *Ccl5, Nkg7, Xcl1, and Ctsw*, demonstrating partial acquisition of a γδT1-like transcriptional state (Fig. S3A-D).

Conversely, analysis of γδT1 populations identified four distinct clusters across the PerC, spleen, and BM, comprising mature cytotoxic γδT1 cells in control PerC and SOX11-induced γδT1 populations with tissue-adapted features in the PerC, a classical γδT1 phenotype in the spleen, and an NK-like phenotype in the BM (Fig. 2F, Fig. S2D-G). Notably, SOX11-induced γδT1 cells also exhibited partial expression of γδT17-associated genes and showed significant overlap with reference γδT17 transcriptional signatures, indicating bidirectional transcriptional plasticity between the γδT17 and γδT1 lineages following SOX11 expression (Fig. S3A-D).

The expansion of γδ T-cells in *SOX11^CD2^* mice was accompanied by a marked reduction in CD4^+^ and CD8^+^ adaptive αβ rearranged T-cell subsets (Fig. 2D-F, Fig. S2F). However, because the *CD2-iCre^tg^* is also active within the B-cell lineage, SOX11 overexpression in *SOX11^CD2^* mice produced prominent alterations in the B-cell development, including a strong expansion of innate B1a cells.

To exclude the possibility that the observed γδ T-cell expansion in *SOX11^CD2^* resulted from indirect effects secondary to altered B-cell development, we crossed *R26-SOX11^tg^* mice with the T-cell restricted *Lck-Cre^tg^* line, yielding *R26-SOX11^tg/tg^; Lck-Cre^tg/+^* mice (hereafter referred to as *SOX11^Lck^*). In this cohort, we also confirmed that SOX11 overexpression significantly increased the proportion of CD4^−^CD8^−^CD3^+^TCRγδ^+^ T-cells in thymus, spleen and bone marrow of *SOX11^Lck^* mice compared to Cre negative control littermates (Fig. 2B,C).

In conclusion, SOX11 overexpression promotes the emergence of both γδT1 and γδT17 subsets across multiple organs, accompanied by a broad shift toward activated type-1/NK-like transcriptional programs at the expense of canonical γδT17 states and a concomitant systemic reduction in adaptive lymphocyte populations.

### SOX11 synergizes with LMO2 to promote γδ-like T-ALL

Next, we investigated whether T-cell-specific SOX11 overexpression is sufficient to drive γδ-like T-ALL in mice, or whether leukemogenesis requires cooperating secondary genetic events. A cohort if 15 *SOX11^Lck^* mice was monitored for up to 500 days, but none of the SOX11 expressing animals developed T-ALL (Fig. 3A,B). This demonstrates that SOX11 overexpression alone is insufficient for malignant transformation of T-cell progenitors and likely requires additional oncogenic hits.

**Figure 3.**
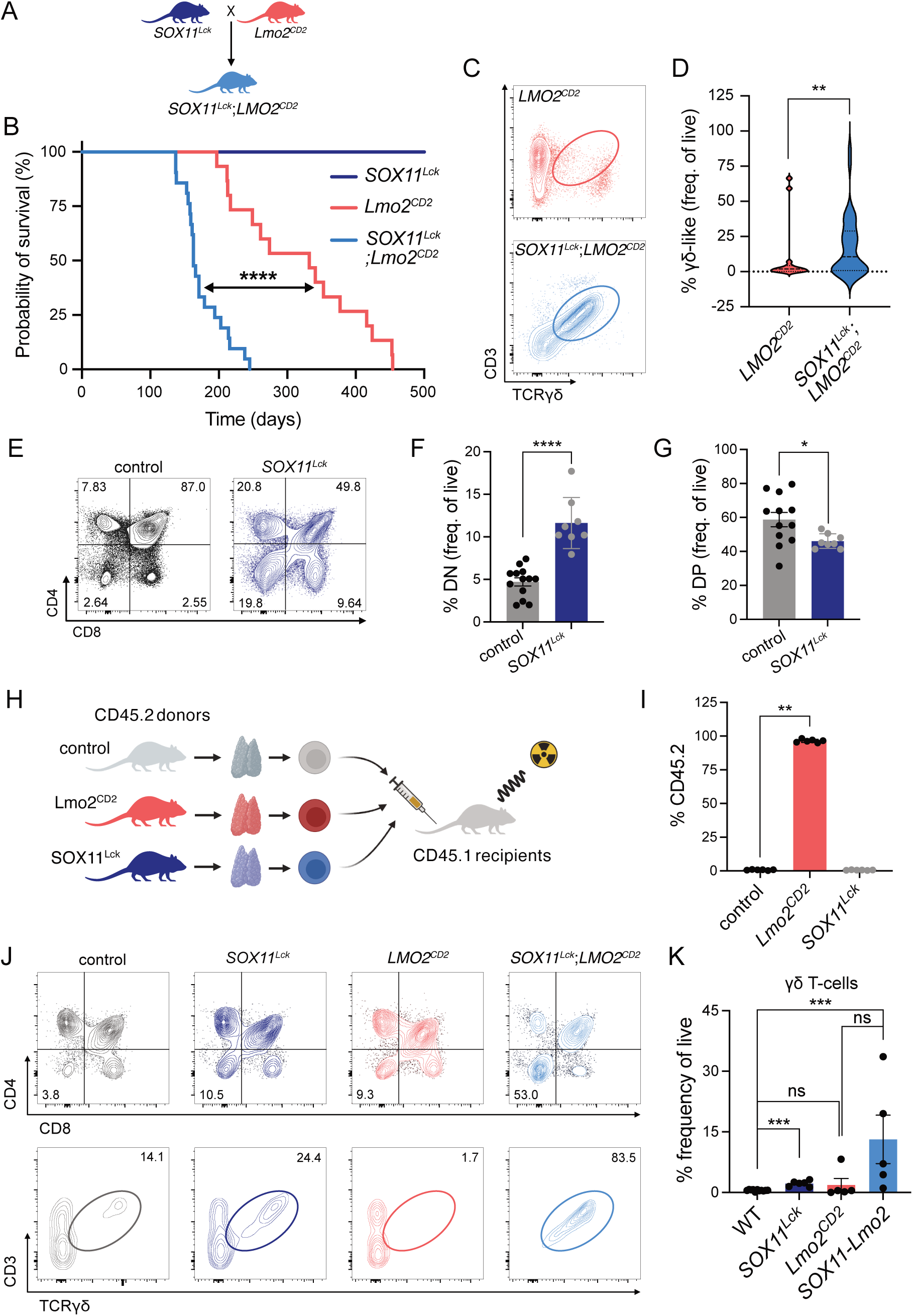
SOX11 synergizes with LMO2 to promote γδ thymocyte development and γδ-like T-ALL. **(A)** Scheme depicting the breeding strategy to generate *SOX11^tg/tg^;LckCre^tg/+^;CD2-Lmo2^tg/+^* (hereafter referred to as *SOX11^Lck^;Lmo2^CD2^*). **(B)** Kaplan Meier survival curve of *SOX11^Lck^* (in dark blue, n = 18), *Lmo2^CD2^* (in red, n = 15), and *SOX11^Lck^;Lmo2^CD2^* mice (in light blue, n = 21). A log-rank Mantel-Cox test showed a statistically significant difference between the survival of *Lmo2^CD2^* and *SOX11^Lck^;Lmo2^CD2^* mice. ****, *p* < 0.0001. **(C)** Representative flow cytometric analysis for CD3 and TCRγδ expression in T-cell leukemias derived from *Lmo2^CD2^* or *SOX11^Lck^; Lmo2^CD2^* mice. Cells were pre-gated on single, live CD90.2^+^CD4^−^CD8^−^ cells. **(D)** Violin plots showing the frequency of CD3^+^TCRγδ^+^ leukemic cells, expressed as a percentage of live cells, in *Lmo2^CD2^* and *SOX11^Lck^;Lmo2^CD2^* mice. CD3^+^TCRγδ^+^ leukemias were identified in 2/15 (13%) *Lmo2^CD2^* tumors and 21/37 (57%) *SOX11^Lck^;Lmo2^CD2^* tumors. A two-sided Fisher’s exact test revealed significant difference between the groups (**; *P* = 0.006). **(E)** Flow cytometric analysis of CD4 and CD8 expression in thymocytes of 8-week-old from mice with T-cell-restricted (*R26-SOX11^tg/tg^;LckCre^tg/+^* hereafter referred to as *SOX11^Lck^*) SOX11 overexpression, or from Cre-negative littermate controls. Cells were pregated for single, live CD90.2^+^ cells. **(F-G)** Bar graphs quantifying the flow cytometry data shown in (A), depicting frequencies of double-negative (DN; CD4^−^CD8^−^; panel F) and double positive (DP; CD4^+^CD8^+^; panel G) thymocyte populations, expressed as a percentage of live cells. **(H)** Diagram showing transplantation of CD45.2+ donor thymocytes from 8-week-old Cre-negative littermate controls or from mice that overexpress SOX11 (*SOX11^Lck^*) or LMO2 (*LMO2^CD2^*) into sublethally irradiated (5.5 Gy) CD45.1 recipients. Six weeks after transplantation, recipient mice were sacrificed and thymic reconstitution was analysed by flow cytometry to determine repopulation by CD45.1^+^ cells derived from the recipient bone marrow or by CD45.2^+^, cells originating from self-renewing donor thymocytes. **(I)** Bar graph showing the percentage of thymocyte recolonization by CD45.2+ donor thymocytes. Whereas thymocytes from *Lmo2^CD2^* mice (n = 6), thymocytes from wild-type (WT; n = 5) or *SOX11^Lck^* (n = 6) mice failed to repopulate the thymus of sublethally irradiated recipients. **(J)** Flow cytometric analysis of thymocytes from 8-week-old Cre-negative littermate controls or from mice expressing SOX11 alone *(SOX11^Lck^)* or Lmo2 alone (*Lmo2^CD2^*) or the combined transgenes (*SOX11^Lck^; Lmo2^CD2^*). γδ T-cells were defined as CD3^+^TCRγδ^+^ cells within the CD4^−^CD8^−^ DN thymocyte population, pregated on single, live CD90.2^+^ cells. **(K)** Bar graphs displaying frequencies of γδ T-cells (as a percentage of live cells) from flow cytometry data shown in (J). Error bars represent the standard error of the mean. * *P* < 0.05, ** *P* <0.01, *** *P* < 0.001, **** *P* <0.0001

Because the high-risk LMO2 γδ-like T-ALL subtype is enriched for genetic alterations leading to LMO2 activation ^10^, we next tested whether SOX11 synergizes with LMO2 during murine T-ALL initiation. *SOX11^Lck^* mice were crossed with the previously described LMO2 overexpression T-ALL mouse models, *CD2-LMO2^tg^* mice ^31^ (hereafter referred to as *LMO2^CD2^*), to generate animals co-expressing both transcription factors (Fig. 3A). Cohorts of *R26-SOX11^tg/tg^; Lck-Cre^tg/+^*; *CD2*-*LMO2^tg/+^* (hereafter referred to as *SOX11^Lck^LMO2^CD2^*) and Cre-negative littermates *R26-SOX11^tg/tg^; Lck-Cre^+/+^;CD2-LMO2^tg/+^* (hereafter referred to as *LMO2^CD2^*) were monitored for spontaneous T-ALL initiation. Combined SOX11 and LMO2 overexpression markedly accelerated leukemia onset (Fig. 3B). Flow cytometric analysis demonstrated that 6 out 11 (55%) *SOX11^Lck^;Lmo2^CD2^* mice developed leukemias resembling human γδ-T-ALL like (Fig. 3C,D). In contrast, γδ-like T-ALL was observed in only one out of seven *Lmo2^CD2^* mice, suggesting that SOX11 cooperates with LMO2 to markedly enhance the development of γδ-like T-ALL in mice, thereby more closely recapitulating the human LMO2-driven γδ-like T-ALL.

Interestingly, SOX11 overexpression also accelerated the initiation of non-γδ LMO2-driven T-ALL, indicating a broader cooperativity between both transcription factors (Fig. S4A-C). To determine whether SOX11-induced acceleration in T-ALL initiation dependent specifically on LMO2 or more generally on other oncogenic drivers, *R26-SOX11^tg/tg^; Lck-Cre^tg/+^* mice were additionally crossed to a second spontaneous murine T-ALL mouse model driven by T-cell restricted loss of the tumor suppressor *Pten*. Cohorts of *R26-SOX11^tg/tg^; Lck-Cre^tg/+^; Pten^fl/fl^* (hereafter referred to as *SOX11/Pten^Lck^*) and *R26-SOX11^+/+^; Lck-Cre^tg/+^; Pten^fl/fl^* (hereafter referred to as *Pten^Lck^*) were monitored longitudinally for T-ALL initiation (Fig. S4D). Unlike the *CD2-LMO2^tg^* model, SOX11 significantly delayed leukemia onset in the context of *Pten* loss (Fig. S4E). This observation is consistent with the high *SOX11* expression observed in TAL1 αβ-like T-ALL, a subtype frequently characterized by *PTEN*-inactivating alterations and associated with a more favorable clinical outcome (Fig. S1E) ^10,32^. Nevertheless, 4 out 8 (50%) analyzed *SOX11/Pten^Lck^* mice developed T-ALL with γδ TCR expression, whereas no γδ TCR-positive leukemias were detected in the *Pten^Lck^* cohort (Fig. S4F-H).

### SOX11 does not confer thymocyte self-renewal but synergizes with LMO2 to promote γδ T-cell formation

A key mechanistic distinction between the *LMO2^CD2^* and the *Pten^Lck^* spontaneous murine T-ALL models is that the *LMO2^CD2^* mice exhibit a pre-leukemic clonal expansion of an immature T-cell progenitor population, which is not seen in the *Pten* loss-of-function model ^33^. Prior studies showed that LMO2 overexpression confers self-renewal capacity in the CD4^−^CD8^−^ double negative (DN) thymocytes, enabling their clonal expansion and accumulation of secondary oncogenic hits, eventually leading to the progression to overt leukemia ^31,33–35^.

To explore how SOX11 affects early thymocyte differentiation, detailed flow cytometric analysis was performed on thymi of 8-weeks old *SOX11^Lck^* mice versus littermate controls. eGFP expression, marking SOX11 overexpression was observed from early DN stage, particularly at the CD44^−^CD25^+^ (DN3) stage (Fig. S5A). SOX11 overexpression caused a partial developmental delay of T-cell differentiation, characterized by a reduced proportion of more mature CD4^+^CD8^+^ double positive (DP) thymocytes and a concomitant higher proportion of DN cells (Fig. 3E-G). Given that overexpression of SOX family members has been linked to the acquisition of stem cell-like properties^36^, and that γδ-like T-ALL subtypes are enriched for ETP-like and stem cell-associated transcriptional programs ^7,9^, we hypothesized that *SOX11^Lck^* thymocytes might acquire pre-leukemic self-renewal capacity, similar to *LMO2^CD2^* thymocytes^31^. To test this hypothesis, we performed thymocyte transplantation experiments, transferring CD45.2^+^ donor thymocytes either overexpressing *SOX11* or *LMO2*, or wild-type thymocytes, into sublethally irradiated CD45.1^+^ syngeneic recipient mice (Fig. 3H). Six weeks post-transplantation, neither the Cre-negative (WT) nor the *SOX11^Lck^* CD45.2^+^ thymocytes were able to reconstitute the thymi of CD45.1^+^ recipient mice (Fig. 3I). In contrast, the *CD2-LMO2^tg^* CD45.2^+^ thymocytes reconstituted the thymus of the irradiated recipient mice (Fig. 3I). We conclude that *SOX11* overexpression alone does not induce pre-leukemic thymocyte self-renewal properties. However, flow cytometric analysis indicates that *SOX11* gain significantly expands the pre-leukemic self-renewing DN3 population in *LMO2^CD2^* mice, while simultaneously permitting/stimulating differentiation towards the γδ T-cell lineage (Fig. 3J,K). The expansion and enforced clonal proliferation of this preleukemic population provides a mechanistic explanation for why accelerated leukemogenesis occurs only when *SOX11* overexpression is combined with *LMO2*, and not with *Pten* loss.

### SOX11 and LMO2 cooperate to induce MYCN transcriptional activity in preleukemic thymocytes

To identify downstream regulators of the combined transcriptional activity of SOX11 and LMO2, we performed RNAseq on FACS sorted DN3 thymocytes from control, *SOX11^Lck^*, LMO2*^tg^* and *SOX11^Lck^;LMO2^tg^* mice (Fig. 4A). Although *SOX11^Lck^* DN thymocytes clustered separately from controls, DN3 *LMO2^CD2^* T-cells cluster together irrespective of *SOX11* expression (Fig. 4B). Relative to control DN3, the most differentially expressed genes were seen in *SOX11^Lck^*, with 2162 genes up- or down-regulated genes. These were highly enriched for MYC target genes (p-adj = 2.71 × 10^−52^, combined score = 393.93; Fig. S6A), as well as genes encoding ribosomal proteins and regulators of translation (p-adj = 3.11 × 10^−71^, combined score = 5051.98). Genes upregulated in *SOX11^Lck^LMO2^CD2^* DN3 were enriched for signatures associated with MYCN targets in high-risk neuroblastoma (p-adj = 1.25×10^−31^, combined score = 804.55; Fig. 4C) and in cell cycle regulation (p-adj = 6.14 × 10^−11^, combined score = 1019.14; Fig. 4D). Notably, MYCN is highly expressed in LMO2 γδ-like T-ALL cases (Fig. 1A; Fig. S6B). Furthermore, a recurrent MYCN P44L mutation, which disrupts a degron motif and is therefore predicted to increase MYCN protein stability, occurs more frequently in SOX11^high^ T-ALL (12/401; 3.0%) compared with SOX11^low^ T-ALL (8/908; 0.9%)(Fig. 4E, Fig. S6C). When T-ALL patients are stratified by molecular subtypes, P44L MYCN mutations occur most frequently in LMO2 γδ-like T-ALL cases (30%; Fig. 4F; Fig. S6D). Notably, two of the three *MYCN* P44L cases were present in the SOX11^high^ group and experienced early clinical events (Fig. S6E,F). Together, these genetic and expression data implicate MYCN as a key driver in the LMO2 γδ-like T-ALL subtype.

**Figure 4.**
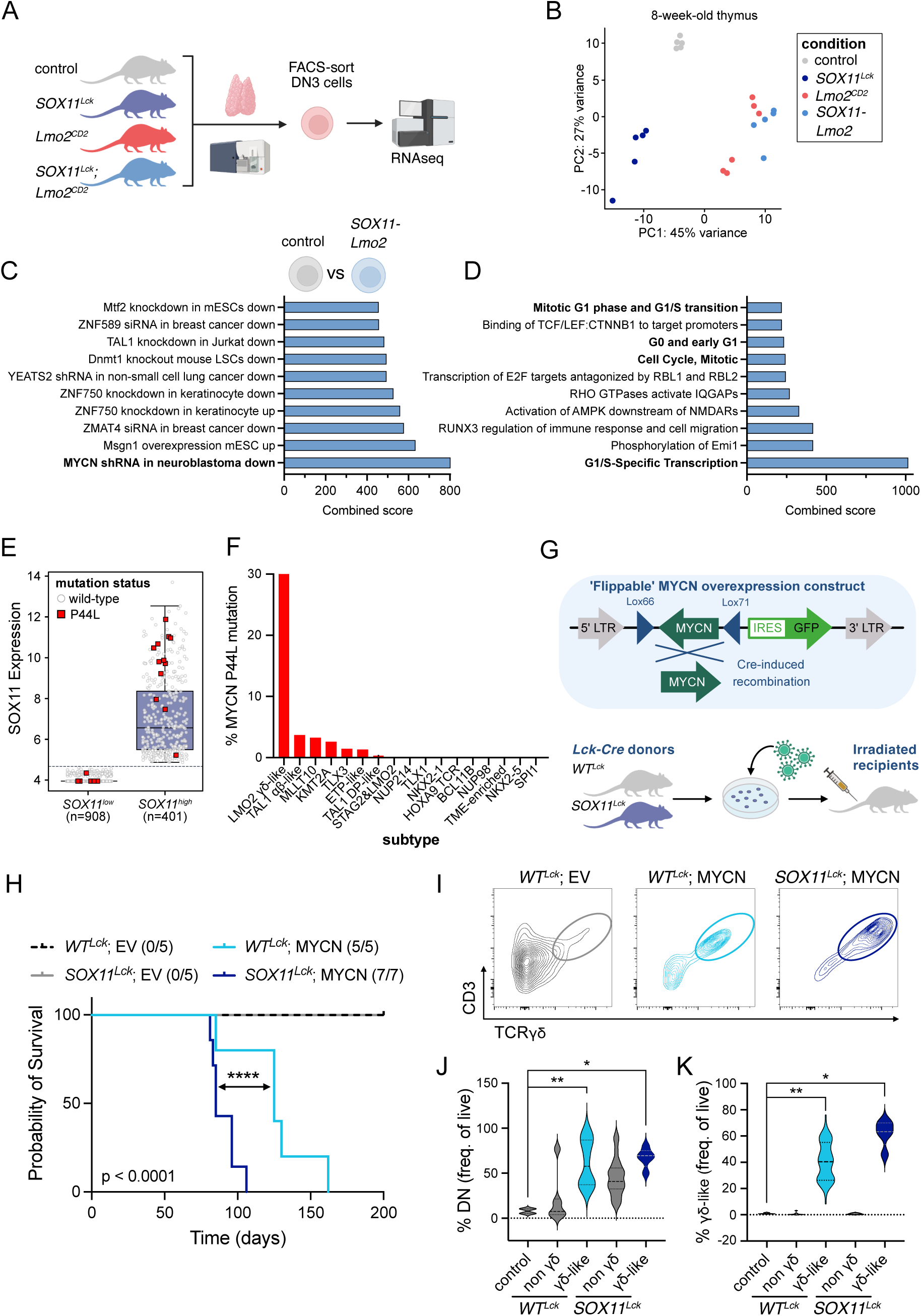
MYCN is a downstream target of the SOX11-LMO2 transcriptional complex, and MYCN cooperates with SOX11 to drive aggressive γδ-like T-ALL. **(A)** Scheme depicting RNAseq analysis of fluorescence-activated cell sorting (FACS)-sorted CD4^−^CD8^−^ double negative (DN3) thymocytes from 8-week-old mice of the indicated genotypes. **(B)** Two-dimensional principal component analysis (PCA) showing genotype-dependent segregation of sorted DN3 thymocytes. Each dot represents a DN thymocyte population derived from an independent mouse. **(C-D)** Bar graphs showing enrichment analysis of genes upregulated in *SOX11^Lck^;Lmo2^CD2^* DN3 thymocytes relative to Cre-negative littermate controls, using the TF Perturbations Followed by Expression (C) or Reactome 2022 Reactome Pathway Database (D). **(E)** Box plots showing the frequency of MYCN P44L mutations in *SOX11^low^* (8/908; 0,9%) and *SOX11^high^* (12/401; 3.0%) T-ALL cases from the Pölönen et al. cohort ^10^. *SOX11^high^* was defined as VST expression > 4.5. Significance was assessed using Fisher’s exact test (**, *P* = 0.0065). **(F)** Bar plot showing the percentage of cases harboring MYCN P44L mutations across T-ALL subtypes ^10^. **(G)** Schematic overview of a Cre-inducible, flippable MYCN overexpression retroviral construct (top) and the bone marrow (BM) transplant assay strategy used to restrict *MYCN* expression to the T-cell lineage. Figure adapted from ^37^. **(H)** Kaplan Meier survival analysis of sublethally irradiated C57BL6 mice transplanted with BM cells from *Lck-Cre*-expressing wild-type (*WT^Lck^*) or SOX11 (*SOX11^Lck^*) donors, transduced with either an empty vector (EV) or a Cre-inducble flippable MYCN overexpression vector. Leukemia penetrance for each cohort is indicated in brackets. Five and three recipient mice transplanted with MYCN-tranduced *WT^Lck^* or *SOX11^Lck^* BM donor cells, respectively, were excluded from the analysis due to the absence of GFP signal at the experimental endpoint (day 250). Combined expression of SOX11 and MYCN significantly accelerates leukemia onset compared with MYCN overexpression alone (log-rank Mantel–Cox test; ****, *P* < 0.0001). **(I)** Flow cytometric analysis for CD4, CD8, CD3 and TCRγδ expression in T-cell leukaemia’s from irradiated recipients transplanted with *WT^Lck^* or *SOX11^Lck^* BM cells, transduced with either an EV or a Cre-inducble flippable MYCN overexpression vector. MYCN-driven T-cell leukemias were compared with non-leukemic splenic cells from *WT^Lck^;* EV recipients. **(J-K)** Violin plots displaying frequencies of CD4^−^CD8^−^ double-negative (DN; J) or CD3^+^TCRγδ^+^ (K) of MYCN-driven leukemic cells or EV-transplanted non-leukemic control cells (expressed as a percentage of live cells), pregated on either single, live CD90.2^+^ (J) or CD90.2^+^CD4^−^CD8^−^ (K) cells. Leukemias arising in MYCN-driven recipients with (n=25) or without (n= 20) SOX11 expression were further stratified into γδ-like (n=5 and 9, respectively) and non-γδ subsets (n=23 and 11, respectively).

We next tested whether MYCN cooperates with SOX11 to accelerate γδ-like T-cell leukemia *in vivo*. To this end, *R26-MYCN^tg^* and *R26-SOX11^tg^* mice were crossed with *Lck-Cre^tg^* mice to generate *R26-MYCN^tg/+^;R26-SOX11^tg/+^;LckCre^tg/+^* mice (hereafter referred to as *SOX11/MYCN^Lck^*), enabling combined *MYCN* and *SOX11* overexpression within the T-cell compartment (Fig. S6G). In this heterozygous setting, monoallelic transgene *MYCN* expression from the *Rosa26* promotor was insufficient to induce T-cell transformation on its own. However, combined monoallelic expression of *SOX11* and *MYCN* synergistically induced T-cell leukemia, albeit with a long latency and penetrance of only 28% (9/32; Fig. S6H).

To validate these results in a more robust experiment, we used a previously published retroviral, Cre-inducible MYCN overexpression vector ^37^ (Fig. 4G, top). Bone marrow hematopoietic progenitors from *Lck-Cre^tg^* control mice *(control^Lck^)* or *SOX11^Lck^* donor mice were transduced with either the Cre-inducible *MYCN* overexpression retroviral vector or an empty control vector (EV). Equal numbers of ex vivo transduced cell population were transplanted into sublethally irradiated syngeneic recipients. Longitudinal monitoring of these mice revealed that SOX11 markedly accelerated MYCN-driven T-cell leukemia development (Fig. 4H). Flow cytometry characterization of the resulting leukemias revealed that γδ-like T-ALL developed in a subset of mice expressing MYCN alone (2 out 5 mice) or combined *MYCN* and *SOX11* (1 out of 7 mice) (Fig. 4I,J). Collectively, these data identify MYCN as a driver of γδ-like T-ALL, with SOX11 acting as a cooperative factor that promotes aggressive disease and accelerates leukemia onset, mirroring the clinical behavior observed in LMO2 γδ-like T-ALL patients.

## Discussion

The outcome of pediatric T-cell acute lymphoblastic leukemia (T-ALL) has improved substantially over recent decades, largely due to intensification of chemotherapy-based treatment regimens. Nevertheless, recent molecular classification studies have demonstrated that T-ALL comprises at least 15 biologically distinct subtypes defined by characteristic gene expression programs and genomic alterations ^10^. Despite this heterogeneity, all T-ALL subtypes, with the exception of ETP-ALL, continue to receive largely identical treatment. These findings highlight the need for subtype-specific therapeutic approaches that exploit the unique molecular dependencies of individual T-ALL entities.

Such an approach may be particularly relevant for patients with SPI1 fusion-positive T-ALL and LMO2 γδ-like T-ALL, two subtypes associated with exceptionally poor clinical outcome ^10^. Among these, LMO2 γδ-like T-ALL is characterized by primary induction failure, rapid disease progression, and early clinical events that frequently result in fatal disease. This subtype appears to originate from unconventional γδ T-cells, a lymphocyte population that combines innate and adaptive immune functions through expression of a rearranged TCRγδ receptor ^29^. Functionally, γδ T-cells in mice can be broadly divided into IL-17-producing γδT17 cells and IFNγ-producing γδT1 cells. Whereas γδT1 cells primarily circulate through blood and lymphoid organs and contribute to anti-viral and anti-tumor immunity, γδT17 cells reside predominantly in barrier tissues and mediate early inflammatory responses.

Historically, T-ALLs expressing a TCRγδ receptor were grouped together as γδ T-ALL and assigned a common prognosis. However, clinical studies yielded conflicting results, with some reporting inferior outcome ^7,8^ and others finding no prognostic impact ^6,9^. More recent genomic analyses have resolved this apparent discrepancy by revealing substantial biological heterogeneity within γδ T-ALL. In particular, pediatric γδ T-ALL diagnosed before three years of age is highly enriched for *LMO2* activation and *STAG2* loss and exhibits an extremely poor prognosis ^7^. Furthermore, recent work demonstrated that HD^+^ transcription factors such as HOXA9 and TLX3 can drive cortical-like γδ T-ALLs that respond relatively well to initial therapy ^9^. In contrast, HD^−^ γδ T-ALLs resemble bona fide γδ T-cells, display ETP-like transcriptional features, and share induction failure with LMO2 γδ-like T-ALL. Importantly, however, these patients do not experience the same reduction in overall survival, further emphasizing that the biological drivers underlying individual γδ T-ALL subtypes, rather than γδ lineage identity itself, determine clinical behavior.

To identify transcriptional regulators associated with LMO2 γδ-like T-ALL, we examined genes selectively enriched in this subtype and identified SOX13 and SOX11. SOX13 is a well-established master regulator of γδT17-cell differentiation and function ^27,38^. In contrast, the role of SOX11 in T-cell biology has remained largely unexplored. Notably, SOX11 and SOX13 are evolutionarily members of distinct SOX family subgroups, despite sharing an HMG-box DNA-binding domain ^39^. Our data demonstrate that SOX11 promotes expansion of γδ T-cells while simultaneously inducing a transcriptional state that combines features of both γδT17 and γδT1 lineages. SOX11-induced γδT17 cells partially lost canonical γδT17 markers, including *Sox13*, *Rorc* and *Il23r*, while acquiring γδT1- and NK-associated transcriptional programs. Conversely, SOX11-induced γδT1 populations retained elements of the γδT17 transcriptional signature. These findings suggest that SOX11 promotes a state of lineage plasticity rather than enforcing commitment toward a single γδ T-cell fate.

Plasticity between γδT17 and γδT1 cells has previously been described during infection, in which γδT17 cells can acquire IFNγ-producing characteristics in response to inflammatory stimuli ^40^. Our findings raise the possibility that SOX11 functions as a transcriptional regulator of this plastic state. Whether SOX11 actively drives lineage conversion or instead maintains cells in an incompletely differentiated intermediate state remains to be determined. Nevertheless, the expansion of these plastic γδ T-cell populations may provide a cellular substrate particularly susceptible to malignant transformation and could therefore contribute to the cell-of-origin of LMO2 γδ-like T-ALL.

Importantly, SOX11 expression alone is not sufficient to predict clinical outcome. Although highly expressed in LMO2 γδ-like T-ALL, SOX11 is also abundant in other subtypes, including TAL1 αβ-like T-ALL, where elevated SOX11 expression is associated with favorable prognosis^10^. Consistent with these clinical observations, SOX11 exerted strikingly different effects in our mouse models depending on the underlying genetic context. SOX11 accelerated leukemogenesis in combination with LMO2 or MYCN, resulting in aggressive γδ-like T-ALL with reduced disease latency. In contrast, SOX11 delayed leukemia development in the context of *Pten* loss. Notably, *PTEN* alterations are common in TAL1-driven T-ALL, suggesting that our mouse models recapitulate the subtype-specific effects observed in patients. Together, these findings indicate that SOX11 acts as a context-dependent modifier of leukemogenesis rather than as a universal oncogenic driver. However, combined *SOX11* and *LMO2* activity is unlikely to fully explain the aggressive biology of LMO2 γδ-like T-ALL. For example, the STAG2-LMO2 subgroup frequently exhibits both high *SOX11* expression and *LMO2* activation but does not consistently display the same poor clinical outcome ^10^. These observations suggest that additional cooperating events are required to drive aggressive leukemia and poor clinical outcomes.

How SOX11 cooperates with LMO2 remains an important question. LMO2 is known to confer stem cell-like properties and induce thymocyte self-renewal ^31^, and bona fide γδ T-cells display transcriptional similarities to stem cell-like ETP populations ^9^. Surprisingly, despite its developmental functions and the established role of the related SOXC family member SOX4 in cellular reprogramming ^41^, SOX11 did not induce self-renewal in T-cell progenitors. Instead, our data suggest that SOX11 primarily acts by expanding the DN3 thymocyte compartment. In the presence of LMO2, this expansion may increase the pool of susceptible pre-malignant cells capable of acquiring additional transforming events. Conversely, by reducing the size of the DP compartment, SOX11 may decrease the number of cells vulnerable to leukemogenesis driven by the loss of PTEN, thereby delaying disease onset ^42^. These observations provide a unifying explanation for the opposing effects of SOX11 in different genetic contexts.

Our data identify MYCN as a strong candidate for this additional cooperating factor. MYCN has previously been implicated as a driver of T-cell lymphomagenesis ^37^ and emerges here as a recurrent feature of SOX11-high disease. In human T-ALL, elevated SOX11 expression is associated with increased MYCN levels, reduced MYC expression, and enrichment of the stabilizing MYCN P44L mutation ^32^. Consistent with these observations, transcriptomic analyses revealed activation of MYC target programs in SOX11-expressing thymocytes, whereas combined SOX11 and LMO2 expression preferentially induced MYCN-associated transcriptional signatures. Furthermore, both MYCN expression and MYCN P44L mutations are particularly enriched in LMO2 γδ-like T-ALL. Most importantly, functional studies demonstrated that SOX11 and MYCN cooperate in vivo to accelerate T-cell leukemogenesis, providing experimental support for a synergistic interaction between these factors. Together, these findings suggest that MYCN activation represents a critical cooperating oncogenic event that contributes to the aggressive clinical behavior of LMO2 γδ-like T-ALL.

Collectively, our findings support a model in which SOX11 promotes expansion and transcriptional plasticity of γδ-lineage progenitors, thereby creating a permissive cellular context for transformation. In isolation, this effect is insufficient to drive aggressive leukemia. However, in the presence of cooperating oncogenic events such as LMO2 and MYCN activation, SOX11 contributes to the development of highly aggressive γδ-like T-ALL. These results identify SOX11 as a key regulator of γδ T-cell biology and provide new insights into the molecular mechanisms underlying one of the most lethal T-ALL subtypes.

## Material and Methods

### Bioinformatic analysis of human T-ALL cohort

Variance-stabilized transformed (VST) RNA-seq expression data and matched clinical, subtype, TCR rearrangement, and mutation annotations for 1309 T-ALL patients were obtained from the Pölönen et al. Synapse dataset (syn54032669)^10^. Analyses were performed in Python 3.12 using pandas, scipy, seaborn/matplotlib, and lifelines. To identify transcription factors enriched in LMO2 γδ-like T-ALL, median expression was calculated for each gene in LMO2 γδ-like cases and compared with the median expression across the remaining T-ALL cohort. Genes were ranked by relative median shift, calculated as:

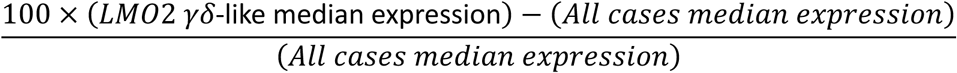

Transcription factors were defined using the curated human transcription factor catalogue from Lambert et al. ^43^. Statistical significance between levels of expression were assessed using a two-sided Mann-Whitney U test. SOX11-positive cases were defined as those with SOX11 VST expression > 4.5, a threshold selected from the cohort-wide SOX11 expression density to distinguish baseline/non-expressing samples from cases with detectable SOX11 expression.

Event-free survival (EFS) was analyzed using Kaplan-Meier estimates. For subtype-specific survival analyses, patients within each subtype were stratified into SOX11-high and SOX11-low groups using a within-subtype median split of SOX11 expression. Survival curves were generated using EFS time and EFS status, and groups were compared using the log-rank test.

Feature-prevalence heatmaps were generated by calculating, within each subtype or classifying-driver group, the percentage of cases positive for selected features, including SOX11 expression > 4.5, TCRG rearrangement, TCRD rearrangement, combined TCRG/TCRD rearrangement, induction failure, and MYCN mutation.

MYCN P44L mutation frequencies were calculated from matched mutation calls. For cohort-wide SOX11-high versus SOX11-low comparisons, patients were stratified using the SOX11 > 4.5 threshold, and enrichment of MYCN P44L mutations was assessed using Fisher’s exact test. MYCN P44L frequency across T-ALL subtypes was calculated as the percentage of cases within each subtype harboring the MYCN P44L mutation.

### H3K27ac HiChIP and ATACseq profiling

Experiments and data preprocessing steps were performed as previously described for 20 published H3K27ac HiChIP and ATACseq samples^10^. Four candidate *SOX11* regulatory elements; chr2:5661022-5662282, chr2:5673202-5674067, chr2:5692079-5692511, chr2:5696261-5697669 were identified using hg38 coordinates in genome paint. As additional descriptive analysis, chromatin signal was quantified from H3K27ac HiChIP and diagnostic ATAC-seq bigWig files as interval-weighted mean signal across each enhancer. *SOX11* z-score derived from vst normalized expression data^10^ was extracted for the corresponding chromatin samples.

### Targeting vector design, mESC targeting and generation of *R26-SOX11^tg^* mice

*R26-SOX11* mice were generated using a previously described optimized Rosa26 recombination-mediated cassette exchange (RMCE)-based targeting strategy ^46,47^. An RMCE-compatible targeting vector (pRMCE-DV1-*SOX11*), containing a floxed transcriptional stop cassette followed by the *SOX11* open reading frame (ORF; NM_003108) and an EGFP reporter, was generated by Gateway cloning. Briefly, the *SOX11* ORF was cloned into the pCR8/GW/TOPO vector (Thermo Fisher Scientific) and subsequently recombined into the pRMCE-DV1 destination vector (BCCM/GeneCorner; LMBP 8870) using Gateway LR recombination. Correct assembly of the targeting construct was confirmed by restriction digest analysis and Sanger sequencing.

G4 ROSALUC mouse embryonic stem cells (mESCs) ^46^ were maintained on mitomycin C-treated mouse embryonic fibroblasts (MEFs; TgN(DR4)1Jae strain) on gelatin-coated plates ^47^. For RMCE, mESCs were co-transfected with pRMCE-DV1-SOX11 and the FlpE recombinase plasmid (pCAGGS-FlpE-IRES-puromycin-pA) using Lipofectamine 2000 (Thermo Fisher Scientific). After 48 h, cells were subjected to G418 selection (200 μg/ml), and resistant colonies were isolated after 7–10 days. Correct Rosa26 targeting was validated by PCR.

Chimeric mice were generated by diploid embryo aggregation at the VIB Transgenic Core Facility (Ghent) as previously described ^47,48^. Briefly, targeted mESCs derived from clone 1 were aggregated with zona pellucida-free E2.5 Swiss embryos and cultured overnight in KSOM medium under mineral oil at 37°C and 5% CO2 before transfer into pseudopregnant B6CBAF1 females. High-grade chimeras were identified by agouti coat color and black eye pigmentation, and germline transmission was established to generate the R26-SOX11 mouse line.

### Murine T-ALL models

Mice with conditional SOX11 overexpression and eGFP reporter were generated according to a previously described optimized *Rosa26* targeting strategy ^46,47^. To induce lymphocyte-specific or T-cell specific SOX11 expression, *R26-SOX11^tg/tg^* mice were crossed with CD2-iCre^49^ (JAX 008520) or Lck-Cre ^50^ animals (JAX 003802), respectively, thus originating the *R26-SOX11^tg/tg^*;*CD2-iCre^tg/+^* (*SOX11^CD2^*) and *R26-SOX11^tg/tg^*;*Lck-Cre^tg/+^* (*SOX11^Lck^*) strains. To produce the murine T-ALL models, *SOX11^Lck^* mice were crossed with either *Pten^fl/fl^*^51^*, CD2-Lmo2^tg/+^* (*Lmo2^CD2^*)^35^, or *R26-Mycn^tg/tg^* mice^52^, to generate *R26-SOX11^tg/tg^*;*Pten^fl/fl^*;*Lck-Cre^tg/+^* (*SOX11/Pten^Lck^*), *R26-SOX11^tg/tg^*;*Lck-Cre^tg/+^*;*CD2-Lmo2^tg/+^* (*SOX11^Lck^;Lmo2^CD2^*), or *R26-SOX11^tg/tg^*;*Mycn^tg/tg^*;*Lck-Cre^tg/+^* (*SOX11/Mycn^Lck^*), respectively. We monitored aging cohorts and plotted their survival using a Kaplan–Meier survival curve while testing for statistically significant differences using the log-rank Mantel–Cox test. All animal experiments were approved by the Animal Experiments Ethics Committee of the University Hospital of Ghent.

### Single-cell RNA and feature-barcode sequencing (CITE-seq)

### Sample preparation and library generation

Freshly isolated peritoneal cavity (PerC), spleen and bone marrow (BM) cells from 16-week-old *SOX11^CD2^* mice (n = 3) and littermate controls (n = 3) were washed with RPMI 1640 medium (Gibco) supplemented with 1% penicillin/streptomycin and 1% L-glutamine (Gibco) and counted. One million live cells per sample were washed with Cell Staining Buffer (BioLegend), blocked with TruStain FcX (BioLegend; 10 min, 4°C) and stained with the TotalSeq™-A Mouse Universal Cocktail v1.0 (BioLegend; 30 min, 4°C, in the dark). Cells were resuspended in buffer containing propidium iodide (Merck Life Science) and live cells were sorted on a BD LSRII or BD FACSFusion (BD Benelux) using a 100-µm nozzle. For each sample, 125,000 viable cells were loaded onto a 10x Genomics Chromium Controller for droplet generation with the Chromium Next GEM Single Cell 5ʹ Kit v2 (10x Genomics). Gene expression (GEX), feature-barcode (FB/antibody-derived tag, ADT) and TCR V(D)J libraries were prepared using the corresponding 10x Genomics Library Construction Kit and Chromium Single Cell Mouse TCR Amplification Kit according to the manufacturer’s instructions and sequenced on an AVITI24 instrument (Element Biosciences).

### Read processing and aggregation

Sequencing reads were processed with the 10x Genomics Cell Ranger pipeline (v9.0.1). Following per-sample quality control, samples from each biological condition (*SOX11^CD2^* and control) were aggregated with sequencing-depth normalization into a common count matrix for each setup (PerC, spleen and BM), yielding paired GEX and antibody-capture (ADT) matrices per organ. Cell Ranger-derived dimensionality reductions, graph-based and k-means clustering, and t-SNE/UMAP projections were inspected in the Loupe Browser (v9.0).

### Downstream single-cell analysis

Downstream analyses were performed in R with the Seurat package (v5.2.1), applying an identical pipeline to each setup (PerC, spleen and BM). For every organ, the aggregated filtered feature-barcode matrix was imported (Read10X); a Seurat object was created from the GEX assay (min.cells = 1, min.features = 100) and the matched antibody-capture counts were added as a separate ADT assay. RNA counts were log-normalized, and the top 3000 variable features were selected and scaled; ADT counts were normalized by per-cell centered log-ratio (CLR) transformation and scaled. Bona fide T cells were identified within each organ using a robust, threshold-based ADT gate that retained CD3^+^ cells positive for TCRβ or TCRγδ, or for auxiliary T-cell markers (CD2, CD5, CD7), and negative for B-cell (CD19/CD20), myeloid (CD14/CD16/CD11b/Ly6G) and NK-cell (CD56/NK1.1) markers; positivity thresholds were set automatically from each marker’s intensity distribution. The gated T-cell compartment was re-processed independently on both modalities: ADT-based clustering used principal component analysis (PCA) over all antibody features (dimensions 1-10, resolution 0.4), and RNA-based clustering used PCA over the variable features (dimensions 1-30, resolution 0.6), each followed by t-SNE and UMAP embedding. Cluster-defining markers were derived with FindAllMarkers, applying an average-expression cut-off of 0.5 during selection of cluster-specific genes.

### Differential expression, functional enrichment and gene-set enrichment analysis

Differential expression between SOX11-overexpressing and control cells was computed within each lineage (γδ T cells and B cells) and each organ using Wilcoxon rank-sum tests (FindMarkers; logfc.threshold = 0, min.pct = 0.05); genes with an adjusted p-value < 0.05 were considered significant, and directional signature gene sets were defined at |log_2_FC| ≥ 0.25. Cross-organ consensus signatures were derived by combining per-organ differentially expressed features. Functional interpretation used clusterProfiler (v4.14.4) with MSigDB gene sets retrieved through msigdbr (Mus musculus; Hallmark and GO:Biological Process collections), applying over-representation analysis (Benjamini-Hochberg-adjusted; enrichment cut-off 0.1) and pre-ranked gene-set enrichment analysis (GSEA; adjusted-p cut-off 0.25). SOX11-induced γδ T-cell target clusters (e.g., the splenic gdT1 and gdT17 populations) were further characterized by differential expression against the remaining reference clusters.

### RNA-extraction, RNA sequencing and RT-qPCR

RNA was extracted using ‘miRNeasy Mini’ (Qiagen) or ‘miRNeasy Micro’ (Qiagen) kits, with on-column DNAase digestion performed according to the manufacturer’s instructions. cDNA synthesis was carried out using iScript™ Advanced cDNA Synthesis Kit (Bio-Rad Laboratories). RT-qPCR was performed using the SsoAdvanced Universal SYBR® Green Supermix (Bio-Rad Laboratories) on a LightCycler 480 system (Roche). Gene expression levels were normalized to reference genes and analyzed using qbase+ software (Biogazelle). Primer sequences are listed in Table S2.

### Immunoblot

Cells were lysed at a density of 20.000 per µL in RIPA buffer supplemented with protease inhibitor cocktail (Roche). The protein content of the lysates was quantified using Pierce BCA Protein Assay (Thermo Fisher), and 12 µg of protein per sample was run on 10% precast polyacrylamide gels (BioRad) at a constant voltage of 100 V for 90 minutes. Afterwards, the protein was transferred onto nitrocellulose membranes (BioRad), and the membrane was incubated with primary antibodies targeting SOX11 (HPA000536; Sigma-Aldrich), Vinculin (clone hVIN-1; Sigma-Aldrich), or β-Actin (clone 2D4H5; Thermo Fisher), and HRP-linked secondary antibodies (Cell Signaling Technology). Images were obtained by chemiluminescent reaction with West Dura substrate (Thermo Fisher), acquired using ChemiDoc (BioRad), processed and analyzed using the ImageLab software (BioRad).

### Flow cytometry

For flow cytometry, 1 × 10^6^ cells were stained at 4°C in the dark with antibodies (Table S3) and measured on FACSymphony A3 (BD) and analyzed using FlowJo software (Tree Star).

### Thymocyte self-renewal

To assess the self-renewal capacity of *SOX11^Lck^* thymocytes, the thymus of 7-10 weeks old mice expressing CD45.2 was collected and a single-cell suspension was made using a 70 µm cell strainer (Falcon). In addition, 8-12 weeks old recipient mice expressing the CD45.1 allele were irradiated with 5.5 Gy to deplete their thymus of lymphoid cells and, 4 hours later, were injected with the donor CD45.2^+^ thymocytes. To prevent infections, in the 2 weeks following irradiation the recipient mice were given 1.7 g/L of neomycin sulfate in water with a pH of 2.5. After 6 weeks, the recipient mice were sacrificed and their thymic composition was analyzed by flow cytometry (BD Biosciences) to determine whether it had been re-colonized by cells expressing CD45.1 (clone A20; Thermo Fisher), coming from the bone marrow of the recipient mouse, or CD45.2 (clone 104; Thermo Fisher), coming from the self-renewing thymocytes of the donor mice. Thymocytes coming from Cre-negative littermates and *Lmo2^CD2^* mice were used as negative and positive control, respectively.

### RNA sequencing of preleukemic and leukemic samples

For RNA sequencing (RNA-seq), thymi from 8-week-old *SOX11^Lck^*, *Lmo2^CD2^*, *SOX11^Lck^;Lmo2^CD2^* or Cre-negative littermate control mice were harvested and processed into single-cell suspensions using 70 µm cell strainers. Cells were stained with antibodies against T-cell surface markers (Table S3), and pre-leukemic CD4^−^CD8^−^CD44^−^CD25^+^ (DN3) and CD4^+^CD8^+^ (DP) thymocyte populations were isolated by fluorescence-activated cell sorting (FACS) using a BD FACSAria Fusion flow cytometer (BD Biosciences). Sorted subsets were lysed in RLT buffer, and the RNA was purified using the RNeasy kit (Qiagen). RNA quality was assessed on a Fragment Analyzer (Agilent), the RNA was converted to cDNA, and the barcoded cDNA libraries were amplified (Lexicon) and analyzed by next-generation sequencing (Illumina). For RNA-seq data analysis, quality control was performed using fastQC and the reads were aligned to the murine genome GRCm38 using Tophat2. DESeq2 was used for differential expression analysis of RNA-seq data.

### Retroviral Cre-inducible MYCN overexpression

Bone marrow transplantation experiments were performed as previously described^37^. Briefly, hematopoietic stem/progenitor cells (HSPCs) were isolated from 6– to 12-week-old in-house bred *SOX11^+/+^;LckCre^tg/+^* (wild-type; *WT^Lck^*), or *SOX11^tg/tg^;LckCre^tg/+^* (*SOX11^Lck^*) donor mice using the EasySep Mouse Hematopoietic Progenitor Cell Isolation Kit (StemCell Technologies). Retroviral particles were generated in HEK293T cells by co-transfection of the MSCV-LoxP-MYCN-LoxP-IRES-GFP expression vector (MYCN) or an empty vector (EV) together with an ecotropic packaging plasmid using JetPEI transfection reagent (Polyplus). Viral supernatants were harvested 48 h and 72 h post-transfection, pooled and filtered.

For retroviral transduction, 1 × 10^6^ HSPCs were cultured in MEM-α medium supplemented with 20% fetal calf serum (Invitrogen), IL-3 (10 ng/ml), IL-6 (10 ng/ml), stem cell factor (SCF; 50 ng/ml), and retronectin (8 μg/ml). Cells were incubated with viral supernatant and transduced by spinfection at 2500 rpm for 90 min at 30 °C. Four hours after transduction, cells were transferred to fresh cytokine-supplemented RPMI 1640 medium. High transduction efficiencies (70–80% GFP^+^ cells) were consistently observed 48 h post-transduction. Subsequently, 1 × 10^6^ GFP^+^ cells were intravenously injected into recipient C57BL/6J mice that had received sublethal irradiation (5.5 Gy) 4–6 h before transplantation. EV-transduced cells were transplanted into five recipient mice, whereas MYCN-transduced cells were transplanted into ten recipient mice. Five and three recipient mice transplanted with MYCN-tranduced *WT^Lck^* or *SOX11^Lck^* BM donor cells, respectively, were excluded from the analysis due to the absence of GFP signal at the experimental endpoint (day 250).

Recipient mice were monitored daily for signs of disease progression, and peripheral blood was analyzed biweekly. Experimental endpoints were defined by > 20% weight loss or by overt signs of illness, including decreased activity, hunched posture, enlarged lymph nodes, or paralysis. Survival analyses were performed using the Kaplan–Meier method.

## Author contributions

AA, PVV, TP and SG: conceived research AA, TP and SG: wrote the paper

AA, IF, KTC, HDW, PP, EVA, ST, and BL: data collection and analysis.

MVB, PN, FS, TT, JC, CM and DT: contributed new reagents, models or analytic tools.

## Acknowledgments

We thank the UGent Core ARTH animal facilities and Core Flow Cytometry for their technical support.

## Bibliography

1. Girardi T, Vicente C, Cools J, De Keersmaecker K. The genetics and molecular biology of T-ALL. Blood. 2017;129(9):1113–1123.

2. Teachey DT, O’Connor D. How I treat newly diagnosed T-cell acute lymphoblastic leukemia and T-cell lymphoblastic lymphoma in children. Blood. 2020;135(3):159–166.

3. Patel AA, Thomas J, Rojek AE, Stock W. Biology and Treatment Paradigms in T Cell Acute Lymphoblastic Leukemia in Older Adolescents and Adults. Curr Treat Options Oncol. 2020;21(7):57.

4. Rheingold SR, Bhojwani D, Ji L, et al. Determinants of survival after first relapse of acute lymphoblastic leukemia: a Children’s Oncology Group study. Leukemia. 2024;38(11):2382–2394.

5. Pui CH, Robison LL, Look AT. Acute lymphoblastic leukaemia. Lancet. 2008;371(9617):1030–1043.

6. Dourthe ME, Courtois L, Andrieu GP, et al. Impact of T-cell Receptor Status on Mutational Landscape and Outcome in T-ALL. Hemasphere. 2023;7(4):e871.

7. Kimura S, Park CS, Montefiori LE, et al. Biologic and Clinical Analysis of Childhood Gamma Delta T-ALL Identifies LMO2/STAG2 Rearrangements as Extremely High Risk. Cancer Discov. 2024;14(10):1838–1859.

8. Pui CH, Pei D, Cheng C, et al. Treatment response and outcome of children with T-cell acute lymphoblastic leukemia expressing the gamma-delta T-cell receptor. Oncoimmunology. 2019;8(8):1599637.

9. Pinton A, Courtois L, Delafoy M, et al. Homeodomain-driven oncogenic diversion to a gammadeltaTCR phenotype in T-cell acute lymphoblastic leukemia. Blood. 2026;147(23):2793–2808.

10. Polonen P, Di Giacomo D, Seffernick AE, et al. The genomic basis of childhood T-lineage acute lymphoblastic leukaemia. Nature. 2024;632(8027):1082–1091.

11. Penzo-Mendez AI. Critical roles for SoxC transcription factors in development and cancer. Int J Biochem Cell Biol. 2010;42(3):425–428.

12. Dy P, Penzo-Mendez A, Wang H, Pedraza CE, Macklin WB, Lefebvre V. The three SoxC proteins--Sox4, Sox11 and Sox12--exhibit overlapping expression patterns and molecular properties. Nucleic Acids Res. 2008;36(9):3101–3117.

13. Dodonova SO, Zhu F, Dienemann C, Taipale J, Cramer P. Nucleosome-bound SOX2 and SOX11 structures elucidate pioneer factor function. Nature. 2020;580(7805):669–672.

14. Qu Y, Zhou C, Zhang J, et al. The metastasis suppressor SOX11 is an independent prognostic factor for improved survival in gastric cancer. Int J Oncol. 2014;44(5):1512–1520.

15. Brennan DJ, Ek S, Doyle E, et al. The transcription factor Sox11 is a prognostic factor for improved recurrence-free survival in epithelial ovarian cancer. Eur J Cancer. 2009;45(8):1510–1517.

16. Decaesteker B, Louwagie A, Loontiens S, et al. SOX11 regulates SWI/SNF complex components as member of the adrenergic neuroblastoma core regulatory circuitry. Nat Commun. 2023;14(1):1267.

17. Grimm D, Bauer J, Wise P, et al. The role of SOX family members in solid tumours and metastasis. Semin Cancer Biol. 2020;67(Pt 1):122–153.

18. de Bont JM, Kros JM, Passier MM, et al. Differential expression and prognostic significance of SOX genes in pediatric medulloblastoma and ependymoma identified by microarray analysis. Neuro Oncol. 2008;10(5):648–660.

19. Liu DT, Peng Z, Han JY, Lin FZ, Bu XM, Xu QX. Clinical and prognostic significance of SOX11 in breast cancer. Asian Pac J Cancer Prev. 2014;15(13):5483–5486.

20. Ek S, Dictor M, Jerkeman M, Jirstrom K, Borrebaeck CA. Nuclear expression of the non B-cell lineage Sox11 transcription factor identifies mantle cell lymphoma. Blood. 2008;111(2):800–805.

21. Mozos A, Royo C, Hartmann E, et al. SOX11 expression is highly specific for mantle cell lymphoma and identifies the cyclin D1-negative subtype. Haematologica. 2009;94(11):1555–1562.

22. Grönroos T, Mäkinen A, Laukkanen S, et al. Clinicopathological features and prognostic value of SOX11 in childhood acute lymphoblastic leukemia. Sci Rep. 2020;10(1):2043.

23. Nasu A, Gion Y, Nishimura Y, et al. Diagnostic Utility of SOX4 Expression in Adult T-Cell Leukemia/Lymphoma. Diagnostics (Basel). 2021;11(5).

24. Kuwahara M, Yamashita M, Shinoda K, et al. The transcription factor Sox4 is a downstream target of signaling by the cytokine TGF-β and suppresses T(H)2 differentiation. Nat Immunol. 2012;13(8):778–786.

25. van de Wetering M, Oosterwegel M, van Norren K, Clevers H. Sox-4, an Sry-like HMG box protein, is a transcriptional activator in lymphocytes. EMBO J. 1993;12(10):3847–3854.

26. Matos DM, Rizzatti EG, Fernandes M, Buccheri V, Falcao RP. Gammadelta and alphabeta T-cell acute lymphoblastic leukemia: comparison of their clinical and immunophenotypic features. Haematologica. 2005;90(2):264–266.

27. Melichar HJ, Narayan K, Der SD, et al. Regulation of gammadelta versus alphabeta T lymphocyte differentiation by the transcription factor SOX13. Science. 2007;315(5809):230–233.

28. Di Giacomo D, Polonen P, Bardelli V, et al. Deregulation of FOXF1/FENDRR from t(14;16)(q32;q24) defines a subtype of high-risk lineage ambiguous leukemia. Blood. 2026.

29. Ribot JC, Lopes N, Silva-Santos B. gammadelta T cells in tissue physiology and surveillance. Nat Rev Immunol. 2021;21(4):221–232.

30. Inacio D, Amado T, Pamplona A, et al. Signature cytokine-associated transcriptome analysis of effector gammadelta T cells identifies subset-specific regulators of peripheral activation. Nat Immunol. 2025;26(3):497–510.

31. McCormack MP, Young LF, Vasudevan S, et al. The Lmo2 oncogene initiates leukemia in mice by inducing thymocyte self-renewal. Science. 2010;327(5967):879–883.

32. Liu Y, Easton J, Shao Y, et al. The genomic landscape of pediatric and young adult T-lineage acute lymphoblastic leukemia. Nat Genet. 2017;49(8):1211–1218.

33. De Coninck S, Roels J, Lintermans B, et al. Tet2 is a tumor suppressor in the pre-leukemic phase of T-cell acute lymphoblastic leukemia. Blood Adv. 2024.

34. Tremblay CS, Curtis DJ. The clonal evolution of leukemic stem cells in T-cell acute lymphoblastic leukemia. Curr Opin Hematol. 2014;21(4):320–325.

35. Smith S, Tripathi R, Goodings C, et al. LIM domain only-2 (LMO2) induces T-cell leukemia by two distinct pathways. PLoS One. 2014;9(1):e85883.

36. Tsang SM, Oliemuller E, Howard BA. Regulatory roles for SOX11 in development, stem cells and cancer. Semin Cancer Biol. 2020;67(Pt 1):3–11.

37. Vanden Bempt M, Debackere K, Demeyer S, et al. Aberrant MYCN expression drives oncogenic hijacking of EZH2 as a transcriptional activator in peripheral T-cell lymphoma. Blood. 2022;140(23):2463–2476.

38. Spidale NA, Sylvia K, Narayan K, et al. Interleukin-17-Producing gammadelta T Cells Originate from SOX13(+) Progenitors that Are Independent of gammadeltaTCR Signaling. Immunity. 2018;49(5):857–872 e855.

39. Kamachi Y, Kondoh H. Sox proteins: regulators of cell fate specification and differentiation. Development. 2013;140(20):4129–4144.

40. Parker ME, Mehta NU, Liao TC, et al. Restriction of innate Tgammadelta17 cell plasticity by an AP-1 regulatory axis. Nat Immunol. 2025;26(8):1299–1314.

41. Takahashi K, Yamanaka S. Induction of pluripotent stem cells from mouse embryonic and adult fibroblast cultures by defined factors. Cell. 2006;126(4):663–676.

42. Xue L, Nolla H, Suzuki A, Mak TW, Winoto A. Normal development is an integral part of tumorigenesis in T cell-specific PTEN-deficient mice. Proc Natl Acad Sci U S A. 2008;105(6):2022–2027.

43. Sun V, Sharpley M, Kaczor-Urbanowicz KE, et al. The Metabolic Landscape of Thymic T Cell Development In Vivo and In Vitro. Front Immunol. 2021;12:716661.

44. Verboom K, Van Loocke W, Volders PJ, et al. A comprehensive inventory of TLX1 controlled long non-coding RNAs in T-cell acute lymphoblastic leukemia through polyA+ and total RNA sequencing. Haematologica. 2018;103(12):e585–e589.

45. Haenebalcke L, Goossens S, Naessens M, et al. Efficient ROSA26-based conditional and/or inducible transgenesis using RMCE-compatible F1 hybrid mouse embryonic stem cells. Stem Cell Rev. 2013;9(6):774–785.

46. Pieters T, T’Sas S, Demoen L, et al. Novel strategy for rapid functional in vivo validation of oncogenic drivers in haematological malignancies. Sci Rep. 2019;9(1):10577.

47. Pieters T, Goossens S, Haenebalcke L, et al. p120 Catenin-Mediated Stabilization of E-Cadherin Is Essential for Primitive Endoderm Specification. PLoS Genet. 2016;12(8):e1006243.

48. de Boer J, Williams A, Skavdis G, et al. Transgenic mice with hematopoietic and lymphoid specific expression of Cre. Eur J Immunol. 2003;33(2):314–325.

49. Hennet T, Hagen FK, Tabak LA, Marth JD. T-cell-specific deletion of a polypeptide N-acetylgalactosaminyl-transferase gene by site-directed recombination. Proc Natl Acad Sci U S A. 1995;92(26):12070–12074.

50. Dankort D, Curley DP, Cartlidge RA, et al. Braf(V600E) cooperates with Pten loss to induce metastatic melanoma. Nat Genet. 2009;41(5):544–552.

51. Althoff K, Beckers A, Bell E, et al. A Cre-conditional MYCN-driven neuroblastoma mouse model as an improved tool for preclinical studies. Oncogene. 2015;34(26):3357–3368.

