## Supplemental data for "SOX11 stimulates γδ T-cell differentiation and synergizes with LMO2 and MYCN to drive γδ-like T-cell acute lymphoblastic leukemia"

#### Supplementary Figures

**Figure S1.** *SOX11* is highly expressed in LMO2 γδ-like T-ALL, a subtype associated with poor clinical outcome.
**Figure S2.** CITE-seq reveals altered γδ T-cell subtype composition in *SOX11$^{CD2}$* mice.
**Figure S3.** SOX11 induces bidirectional transcriptional plasticity between the γδT17 and γδT1 lineages.
**Figure S4.** Role of SOX11 in leukemia survival and promotion of γδ-Like T-ALL in *Lmo2$^{CD2}$* and *Pten$^{Lck}$* mouse models.
**Figure S5.** Monitoring in vivo SOX11 activation and enhanced DN3 thymocyte development following combined SOX11 and LMO2 expression.
**Figure S6.** High SOX11 expression associates with MYCN P44L mutations and poor clinical outcome in LMO2 γδ-like T-ALL, while SOX11 and MYCN cooperate to drive T-ALL in vivo.

#### Supplementary Tables

**Table S1.** Gentoyping primers
**Table S2.** qRT-PCR primers
**Table S3.** List of antibodies used for flow cytometry

#### Supplementary Figures

#### Figure S1

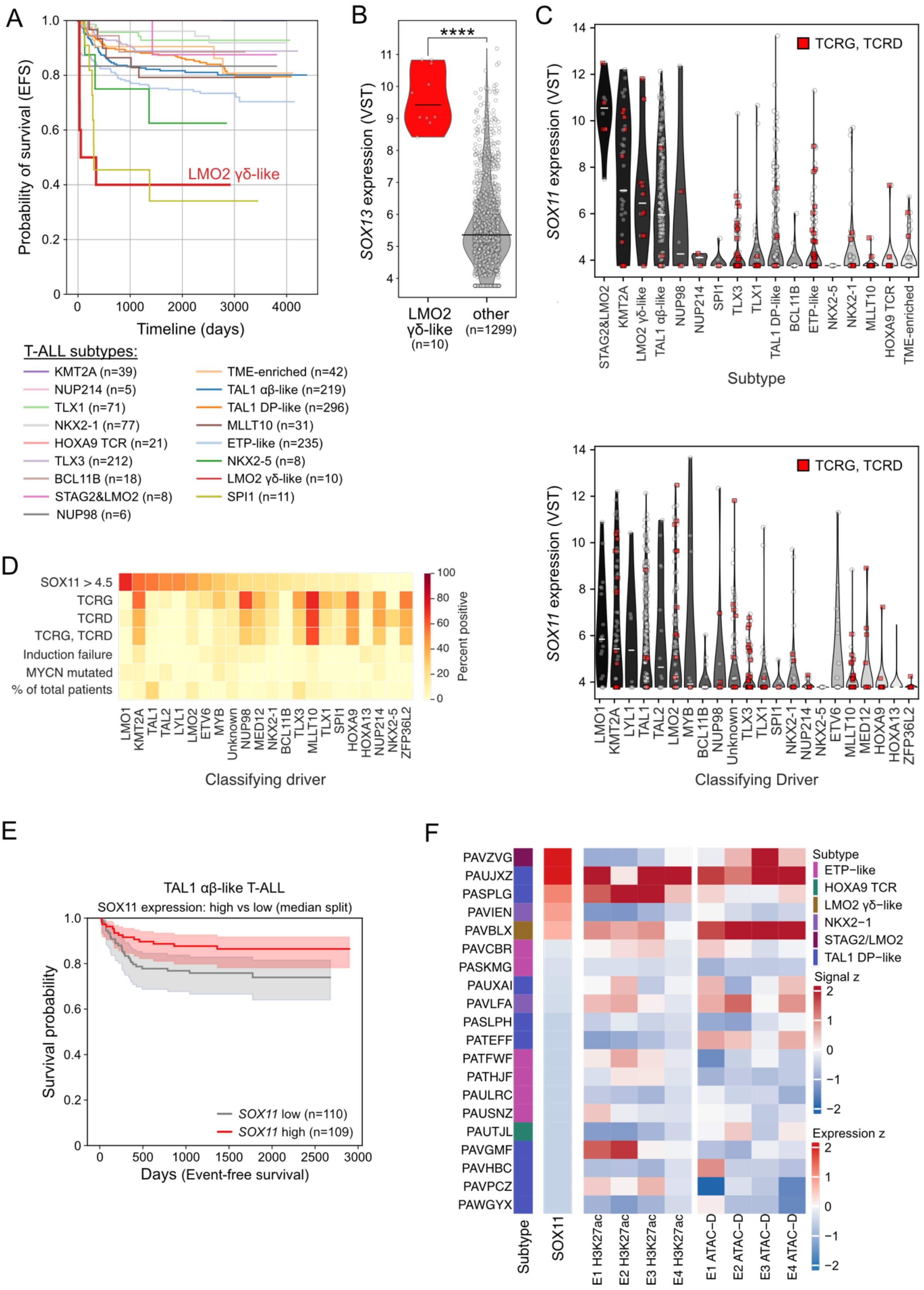

**Figure S1.** ***SOX11*** **is highly expressed in LMO2 γδ-like T-ALL, a subtype associated with poor clinical outcome. (A)** Event-free survival data of 1309 T-ALL patients that are stratified according to their subtype [1]. **(B)** Violin plot depicting *SOX13* expression after variance-stabilizing transformation (VST) in LMO2 γδ-like T-ALL (n=10) compared with all other T-ALL subtypes (n=1293). A Mann-Whitney test showed a significant difference. ****, $P < 0.0001$. **(C)** Violin plot depicting *SOX11* expression of 1309 T-ALL patients [1] that are stratified according to their subtype (top) or classifying driver (bottom). Patients expressing a γδ-TCR, encoded by the TCRG and TCRD loci, are indicated by red squares. **(D)** Heatmap showing the prevalence, plotted as percentage positive, of features (rows) across different classifying drivers (columns). **(E)** Event-free survival of LMO2 γδ-like T-ALL cases from Polonen et al. [1] stratified according to *SOX11* expression (median split). Log-rank analysis revealed a significant difference in outcome between between patients with high versus low SOX11 expression (**, $P = 0.02106$). **(F)** Heatmap showing *SOX11* expression and chromatin signal across four candidate *SOX11* regulatory elements in T-ALL samples. Rows represent samples with matched *SOX11* expression and H3K27ac signal at candidate *SOX11* elements. Samples are ordered by decreasing *SOX11* expression. Columns show T-ALL subtype annotation, *SOX11* expression, H3K27ac signal, and diagnostic ATAC signal across E1 H3K27ac distal, E2 ATAC distal, E3 promoter ATAC, and E4 intragenic/UTR H3K27ac elements. Expression and chromatin values were z-scored by feature across samples and clipped to ±2. Red indicates higher relative expression or chromatin signal, blue indicates lower relative signal. Subtype colors indicate subtype classifications.

**Figure S2**

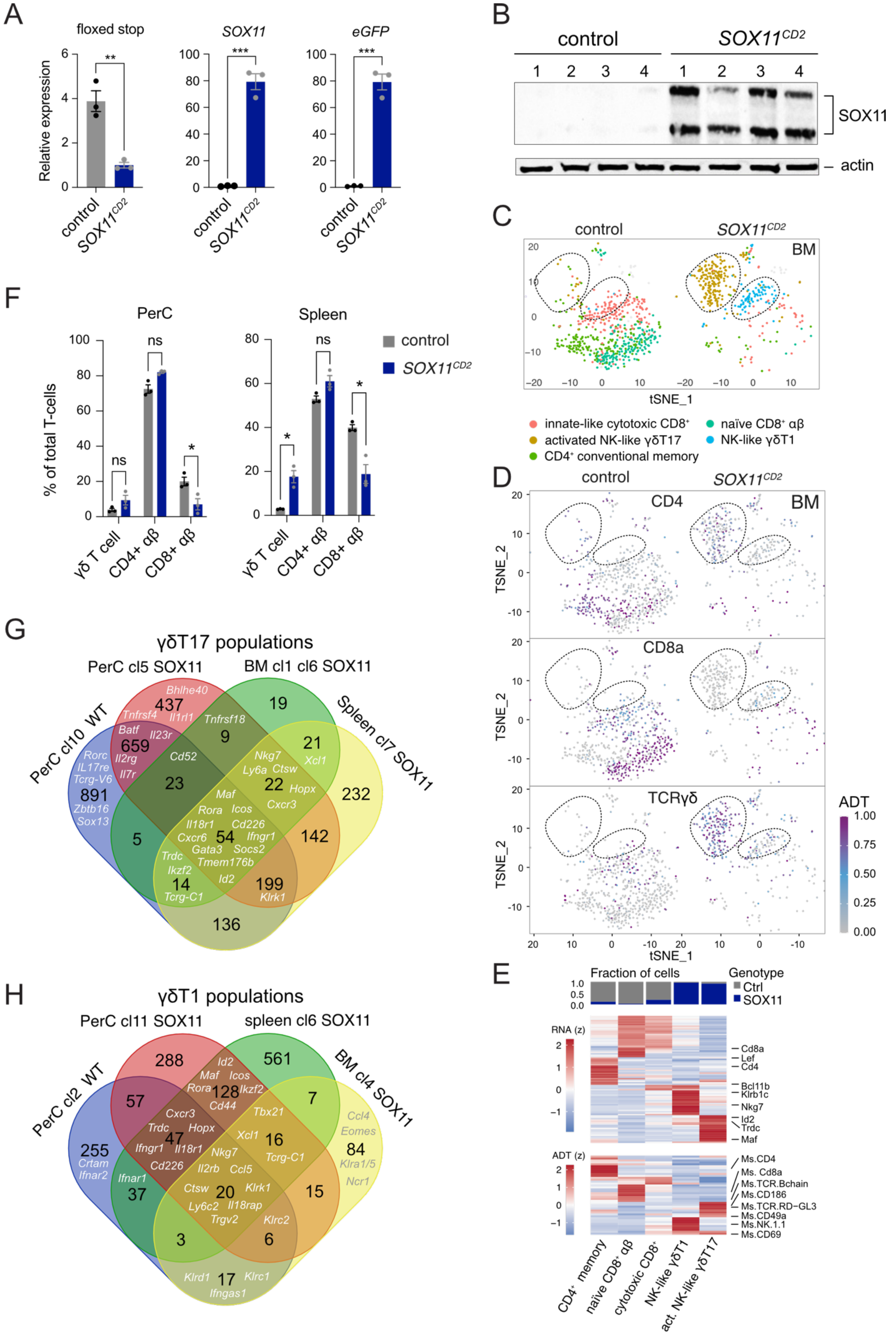
A
floxed stop
SOX11
eGFP
Relative expression
control
SOX11CD2
**
***
B
control
SOX11CD2
1 2 3 4 1 2 3 4
SOX11
actin
C
control
SOX11CD2
BM
tSNE_1
innate-like cytotoxic CD8+
activated NK-like γδT17
CD4+ conventional memory
naïve CD8+ αβ
NK-like γδT1
D
control
SOX11CD2
CD4
CD8a
TCRγδ
BM
TSNE_2
tSNE_1
ADT
1.00
0.75
0.50
0.25
0.00
E
Fraction of cells
Genotype
Ctrl
SOX11
RNA (z)
Cd8a
Lef
Cd4
Bcl11b
Klrb1c
Nkg7
Id2
Trdc
Maf
ADT (z)
Ms.CD4
Ms. Cd8a
Ms.TCR.Bchain
Ms.CD186
Ms.TCR.RD−GL3
Ms.CD49a
Ms.NK.1.1
Ms.CD69
CD4+ memory
naïve CD8+ αβ
cytotoxic CD8+
NK-like γδT1
act. NK-like γδT17
F
PerC
Spleen
control
SOX11CD2
% of total T-cells
γδ T cell
CD4+ αβ
CD8+ αβ
ns
*
G
γδT17 populations
PerC cl5 SOX11
BM cl1 cl6 SOX11
PerC cl10 WT
Spleen cl7 SOX11
437
19
659
9
21
891
23
22
232
5
54
142
14
199
136
Bhlhe40
Tnfrsf4
Il1rl1
Tnfrsf18
Batf
Il23r
Rorc
IL17re
Tcrg-V6
Zbtb16
Sox13
Il2rg
Il7r
Cd52
Nkg7
Ly6a
Ctsw
Xcl1
Hopx
Maf
Cxcr3
Rora
Icos
Il18r1
Cd226
Cxcr6
Ifngr1
Gata3
Socs2
Tmem176b
Trdc
Ikzf2
Id2
Tcrg-C1
Klrk1
H
γδT1 populations
PerC cl11 SOX11
spleen cl6 SOX11
PerC cl2 WT
BM cl4 SOX11
288
561
57
128
7
255
47
16
84
37
20
15
3
6
17
Id2
Maf
Icos
Rora
Ikzf2
Cd44
Cxcr3
Tbx21
Trdc
Hopx
Xcl1
Ifngr1
Il18r1
Nkg7
Tcrg-C1
Ccl4
Eomes
Klra1/5
Ncr1
Crtam
Ifnar2
Cd226
Il2rb
Ccl5
Ifnar1
Ctsw
Klrk1
Ly6c2
Il18rap
Trgv2
Klrc2
Klrd1
Klrc1
Ifngas1

**Figure S2. CITE-seq reveals altered γδ T-cell subtype composition in *SOX11^CD2^* mice. (A-B)** Validation of SOX11 overexpression in splenocytes from 8-week-old control or *SOX11*$^{CD2}$ mice using qRT-PCR (A) or Western blotting (B). **(C)** t-SNE plot showing annotated single-cell clusters from control and SOX11-expressing (*SOX11*$^{CD2}$*)* BM cells. Clusters were defined based on surface marker expression derived from FB-seq. Dotted lines indicate clusters enriched or specifically induced upon SOX11 expression. **(D)** t-SNE plot of single-cell clusters (annotated as in E) from control and SOX11-expressing (*SOX11*$^{CD2}$*)* BM cells, overlaid with cell surface protein expression measured by ScFB-seq for Ms. CD4, Ms. CD8a, and Ms. TCR-γδ (GL3). Dotted lines indicate clusters enriched or specifically induced upon SOX11 expression. **(E)** (top) Bar graph showing, for each BM cluster, the relative proportions of control and SOX11-expressing (*SOX11*$^{CD2}$*)* cells. Heatmaps showing differentially expressed cell surface markers (middle) and genes (middle) in different clusters. ADT: Antibody-determining tag. **(F)** Bar graphs showing the frequency of γδ T-cells, $CD4^+$ αβ cells, and $CD8^+$ αβ T cells, quantified by CITEseq (ADT) and expressed as a percentage of total $CD3^+$ T-cells, in PerC and spleen of control or *SOX11*$^{CD2}$ mice. Statistical significance was determined using an unpaired two-tailed Welch's *t*-test. *p* values: *, $p < 0.05$. ns: not significant. Error bars represent mean ± SEM. **(G-H)** Venn diagram illustrating the overlap of genes expressed among γδT17 (F) and γδT1 (G) clusters from the peritoneal cavity (Perc), spleen, and bone-marrow (BM) of control (WT, wild-type) or *SOX11*$^{CD2}$ mice. Number represent the amount of shared genes between clusters, with selected genes highlighted in white or grey.

**Figure S3**

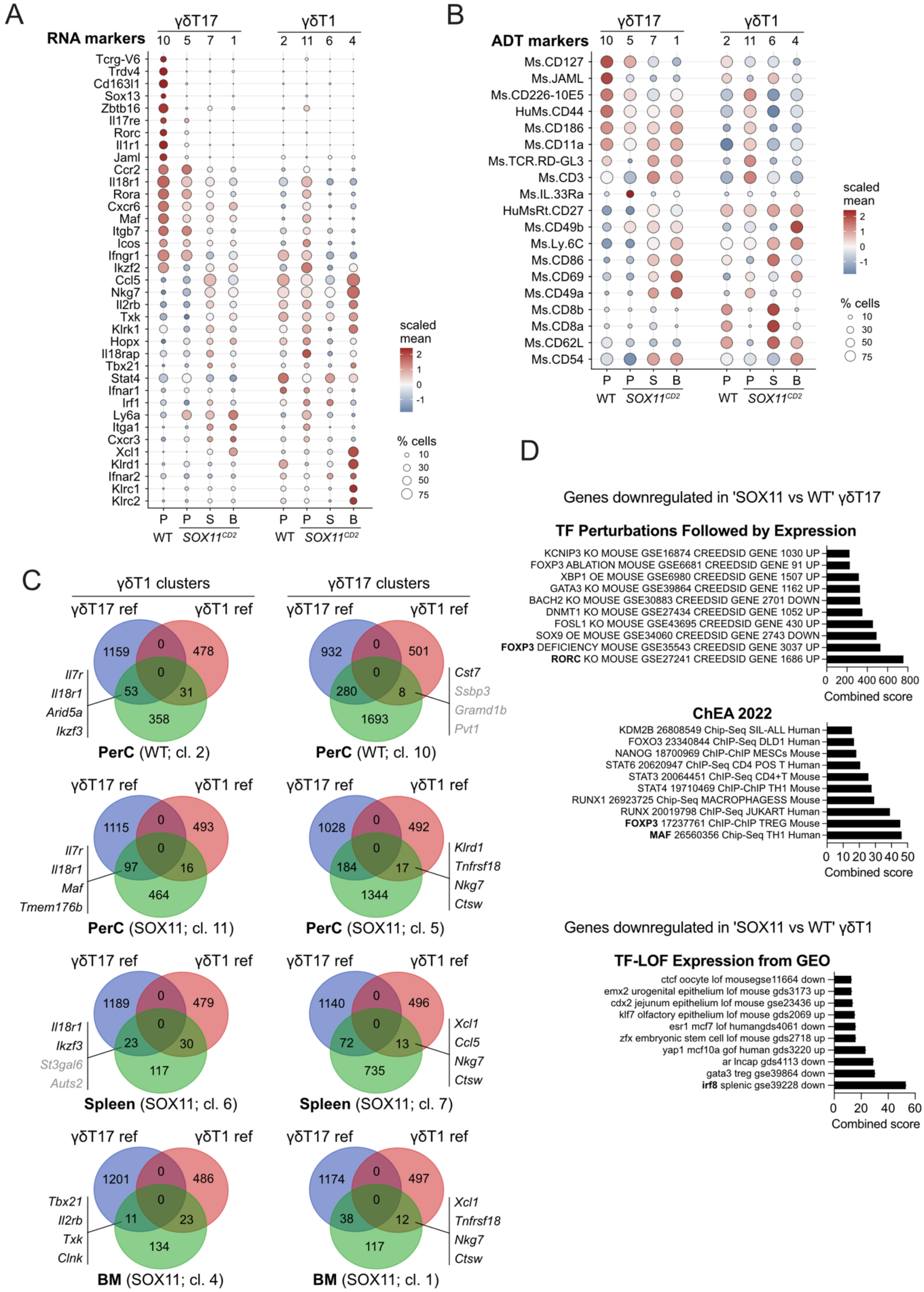

A
RNA markers
γδT17
γδT1
10 5 7 1 2 11 6 4
Tcrg-V6
Trdv4
Cd163l1
Sox13
Zbtb16
Il17re
Rorc
Il1r1
Jaml
Ccr2
Il18r1
Rora
Cxcr6
Maf
Itgb7
Icos
Ifngr1
Ikzf2
Ccl5
Nkg7
Il2rb
Txk
Klrk1
Hopx
Il18rap
Tbx21
Stat4
Ifnar1
Irf1
Ly6a
Itga1
Cxcr3
Xcl1
Klrd1
Ifnar2
Klrc1
Klrc2
P P S B P P S B
WT SOX11CD2 WT SOX11CD2
scaled mean
2 1 0 -1
% cells
10 30 50 75
B
ADT markers
Ms.CD127
Ms.JAML
Ms.CD226-10E5
HuMs.CD44
Ms.CD186
Ms.CD11a
Ms.TCR.RD-GL3
Ms.CD3
Ms.IL.33Ra
HuMsRt.CD27
Ms.CD49b
Ms.Ly.6C
Ms.CD86
Ms.CD69
Ms.CD49a
Ms.CD8b
Ms.CD8a
Ms.CD62L
Ms.CD54
C
γδT1 clusters
γδT17 clusters
γδT17 ref
γδT1 ref
1159 0 478 0 53 31 358
Il7r Il18r1 Arid5a Ikzf3
PerC (WT; cl. 2)
932 0 501 0 280 8 1693
Cst7 Ssbp3 Gramd1b Pvt1
PerC (WT; cl. 10)
1115 0 493 0 97 16 464
Il7r Il18r1 Maf Tmem176b
PerC (SOX11; cl. 11)
1028 0 492 0 184 17 1344
Klrd1 Tnfrsf18 Nkg7 Ctsw
PerC (SOX11; cl. 5)
1189 0 479 0 23 30 117
Il18r1 Ikzf3 St3gal6 Auts2
Spleen (SOX11; cl. 6)
1140 0 496 0 72 13 735
Xcl1 Ccl5 Nkg7 Ctsw
Spleen (SOX11; cl. 7)
1201 0 486 0 11 23 134
Tbx21 Il2rb Txk Clnk
BM (SOX11; cl. 4)
1174 0 497 0 38 12 117
Xcl1 Tnfrsf18 Nkg7 Ctsw
BM (SOX11; cl. 1)
D
Genes downregulated in 'SOX11 vs WT' γδT17
TF Perturbations Followed by Expression
KCNIP3 KO MOUSE GSE16874 CREEDSID GENE 1030 UP
FOXP3 ABLATION MOUSE GSE6681 CREEDSID GENE 91 UP
XBP1 OE MOUSE GSE6980 CREEDSID GENE 1507 UP
GATA3 KO MOUSE GSE39864 CREEDSID GENE 1162 UP
BACH2 KO MOUSE GSE30883 CREEDSID GENE 2701 DOWN
DNMT1 KO MOUSE GSE27434 CREEDSID GENE 1052 UP
FOSL1 KO MOUSE GSE43695 CREEDSID GENE 430 UP
SOX9 OE MOUSE GSE34060 CREEDSID GENE 2743 DOWN
FOXP3 DEFICIENCY MOUSE GSE35543 CREEDSID GENE 3037 UP
RORC KO MOUSE GSE27241 CREEDSID GENE 1686 UP
0 200 400 600 800
Combined score
ChEA 2022
KDM2B 26808549 Chip-Seq SIL-ALL Human
FOXO3 23340844 ChIP-Seq DLD1 Human
NANOG 18700969 ChIP-ChIP MESCs Mouse
STAT6 20620947 ChIP-Seq CD4 POS T Human
STAT3 20064451 ChIP-Seq CD4+T Mouse
STAT4 19710469 ChIP-ChIP TH1 Mouse
RUNX1 26923725 Chip-Seq MACROPHAGESS Mouse
RUNX 20019798 ChIP-Seq JUKART Human
FOXP3 17237761 ChIP-ChIP TREG Mouse
MAF 26560356 Chip-Seq TH1 Human
0 10 20 30 40 50
Combined score
Genes downregulated in 'SOX11 vs WT' γδT1
TF-LOF Expression from GEO
ctcf oocyte lof mousegse11664 down
emx2 urogenital epithelium lof mouse gds3173 up
cdx2 jejunum epithelium lof mouse gse23436 up
klf7 olfactory epithelium lof mouse gds2069 up
esr1 mcf7 lof humangds4061 down
zfx embryonic stem cell lof mouse gds2718 up
yap1 mcf10a gof human gds3220 up
ar lncap gds4113 down
gata3 treg gse39864 down
irf8 splenic gse39228 down
0 20 40 60
Combined score

**Figure S3. SOX11 induces bidirectional transcriptional plasticity between the γδT17 and γδT1 lineages (A-B)** Dot plots showing cell-surface markers (A) or genes (B) that are upregulated (red) or downregulated (blue) in γδT1 and γδT17 subsets across different organs, including peritoneal cavity (PerC, P), spleen (S), and bone marrow (BM, B), from either control or *SOX11*$^{CD2}$ mice. The circle size represents the percentage of γδ cells that are positive for that marker. **(C)** Venn diagram showing the overlap between transcriptional signature of γδT1 and γδT17 subsets from control or *SOX11*$^{CD2}$ mice and published reference γδT1 and γδT17 gene signatures ($P < 0.05$, fold change > 2) [2] . **(D)** Gene set enrichment analysis using genes downregulated upon SOX11 overexpression in PerC γδT-cell subsets relative to controls.

**Figure S4**

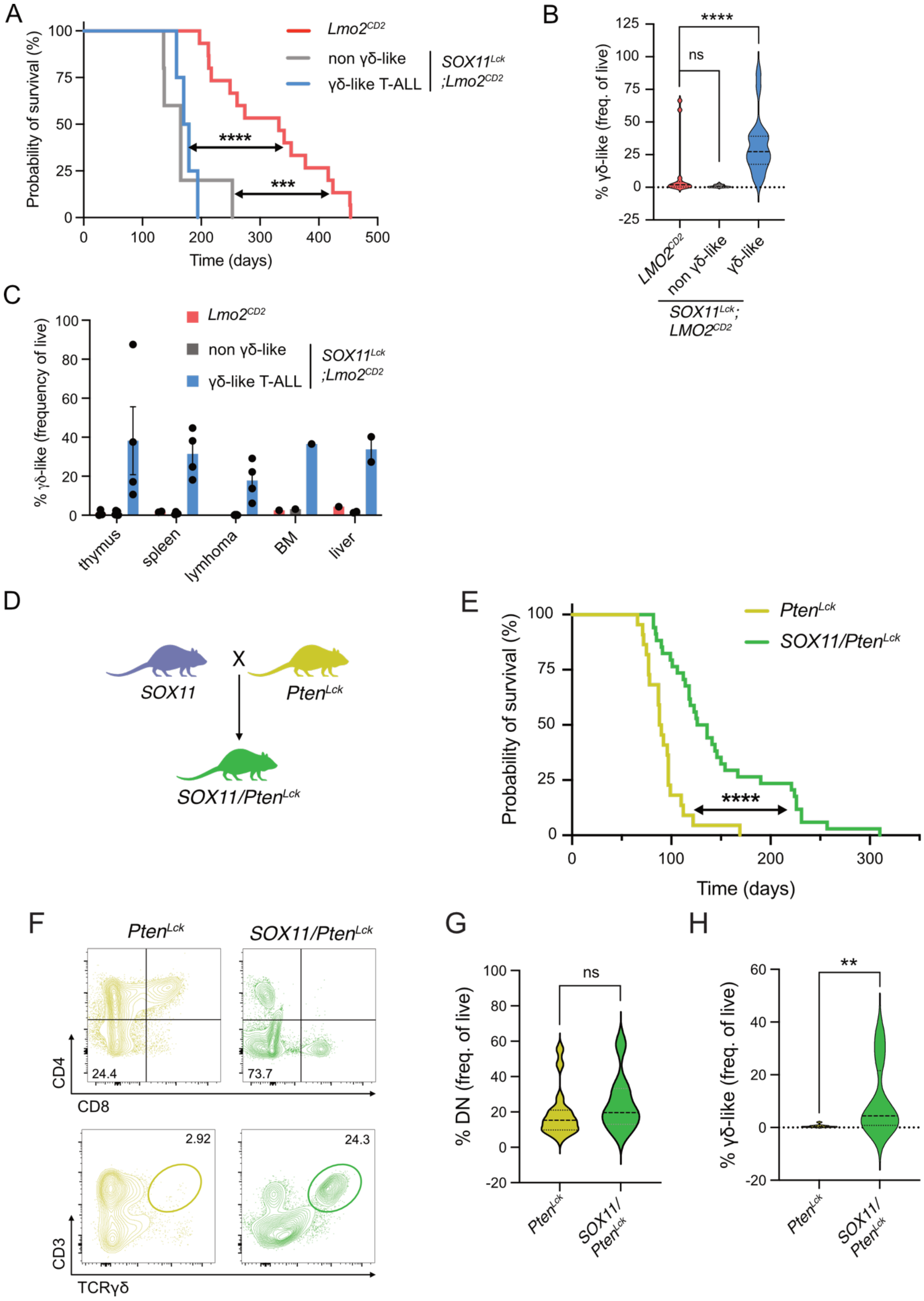
A
Probability of survival (%)
Time (days)
Lmo2CD2
non γδ-like
γδ-like T-ALL
SOX11Lck ;Lmo2CD2
****
***
B
% γδ-like (freq. of live)
****
ns
LMO2CD2
non γδ-like
γδ-like
SOX11Lck; LMO2CD2
C
% γδ-like (frequency of live)
Lmo2CD2
non γδ-like
γδ-like T-ALL
SOX11Lck ;Lmo2CD2
thymus
spleen
lymhoma
BM
liver
D
SOX11
X
PtenLck
SOX11/PtenLck
E
Probability of survival (%)
Time (days)
PtenLck
SOX11/PtenLck
****
F
PtenLck
SOX11/PtenLck
24.4
73.7
CD4
CD8
2.92
24.3
CD3
TCRγδ
G
% DN (freq. of live)
ns
PtenLck
SOX11/ PtenLck
H
% γδ-like (freq. of live)
**
PtenLck
SOX11/ PtenLck

**Figure S4. Role of SOX11 in leukemia survival and promotion of γδ-Like T-ALL in *Lmo2*$^{CD2}$ and *Pten*$^{Lck}$ mouse models. (A)** Kaplan Meier survival curve of *Lmo2*$^{CD2}$ mice (red, n = 15) and *SOX11*$^{Lck}$*;Lmo2*$^{CD2}$ mice developing either γδ-like (blue; n =4) or non-γδ-like (grey; n = 5) leukemias. Survival differences were assessed using a log-rank (Mantel–Cox) test, revealing significantly reduced survival of both γδ-like (**** $P < 0.0001$) and non-γδ-like (*** $P = 0.0002$) *SOX11*$^{Lck}$*;Lmo2*$^{CD2}$ cohorts compared to *Lmo2*$^{CD2}$ mice. **(B)** Violin plots showing the frequency of $CD3^{+}TCR\gamma\delta^{+}$ leukemic cells, expressed as a percentage of live cells, in *Lmo2*$^{CD2}$ and *SOX11*$^{Lck}$*;Lmo2*$^{CD2}$ mice. *SOX11*$^{Lck}$*;Lmo2*$^{CD2}$ tumors were segregated in non γδ-like (n=16) and γδ-like (n=21) tumors and compared to 15 *Lmo2*$^{CD2}$ tumors. A Mann-Whitney test revealed a significant difference between the *Lmo2*$^{CD2}$ and γδ-like groups (**** $P < 0.0001$), whereas no significant difference was observed between the *Lmo2*$^{CD2}$ and non-γδ-like groups (ns; $P = 0.0802$). **(C)** Bar graphs showing the frequencies of γδ-like T-cell leukemias (as a percentage of live cells) across different tumor sites (organs). Data are shown for *Lmo2*$^{CD2}$ and *SOX11*$^{Lck}$*;Lmo2*$^{CD2}$ mice, stratified into γδ-like and non-γδ-like subsets. **(D)** Scheme illustrating the breeding strategy used to generate *SOX11*$^{tg/tg}$*;Pten*$^{fl/fl}$*;LckCre*$^{tg/+}$ (hereafter referred to as *SOX11/Pten*$^{Lck}$) by crossing R26-SOX11tg/tg mice with T-cell-restricted Pten knock out mice (*Pten*$^{fl/fl}$*;LckCre*$^{tg/+}$ hereafter referred to as *Pten*$^{Lck}$). **(E)** Kaplan Meier survival curve of *Pten*$^{Lck}$ (yellow, n = 22), and *SOX11/Pten*$^{Lck}$ mice (green, n = 34). A log-rank Mantel-Cox test showed a statistically significant difference between the survival of PtenLck and SOX11/PtenLck mice. **** $P < 0.0001$. **(F)** Flow cytometric analysis for CD4, CD8, CD3 and TCRγδ expression in T-cell leukaemia's from PtenLck or SOX11/PtenLck mice. **(G-H)** Violin plot displaying frequencies of CD4-CD8- double-negative (DN; G) or CD3+TCRγδ+ (H) leukemic cells (expressed as a percentage of live cells) in PtenLck (n=17) or SOX11/PtenLck (n=8) leukemias, pregated on either single, live CD90.2+ (G) or CD90.2+CD4-CD8- (H) cells. Leukemias arising in *SOX11/Pten*$^{Lck}$ mice were further stratified into γδ-like (n=4) and non-γδ-like (n=4) subsets.

**Figure S5**

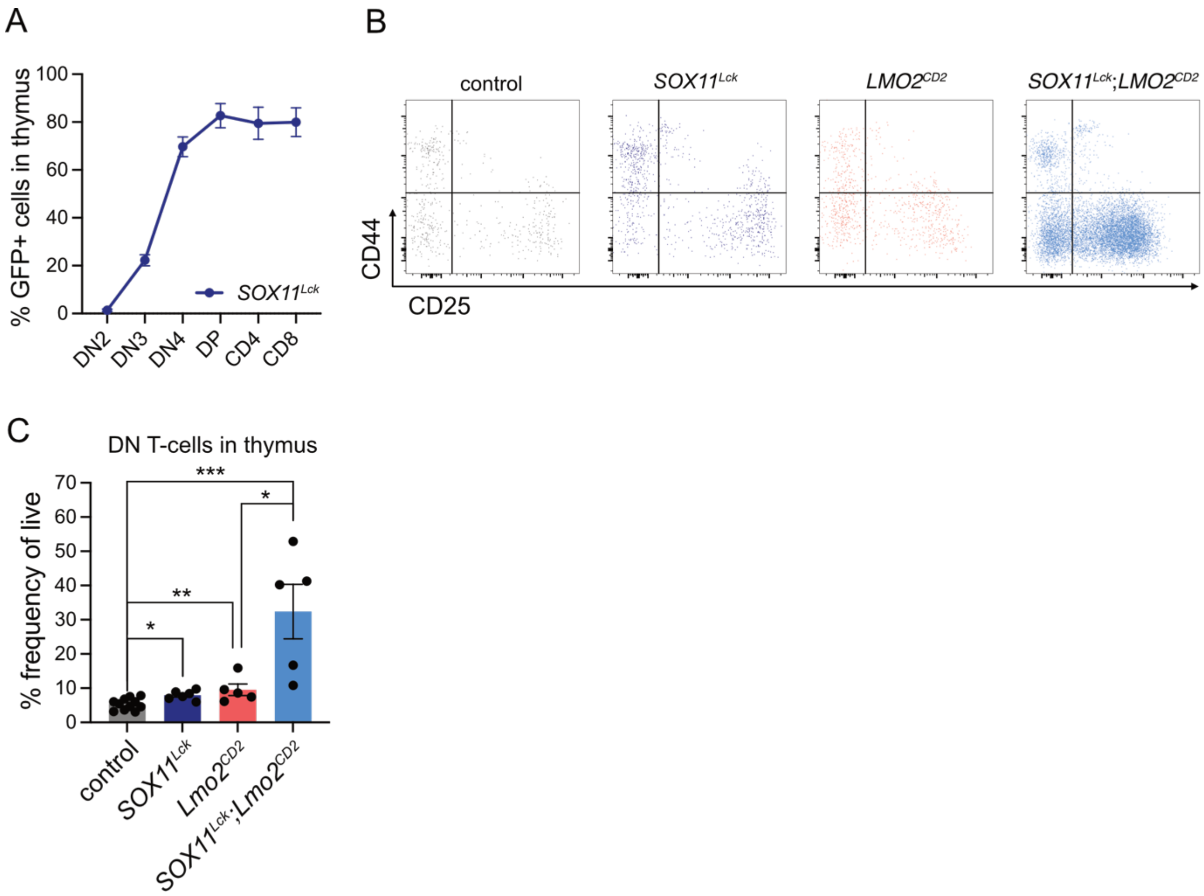
A
% GFP+ cells in thymus
100
80
60
40
20
0
SOX11^Lck
DN2
DN3
DN4
DP
CD4
CD8
B
control
SOX11^Lck
LMO2^CD2
SOX11^Lck;LMO2^CD2
CD44
CD25
C
DN T-cells in thymus
% frequency of live
70
60
50
40
30
20
10
0
***
*
**
*
control
SOX11^Lck
Lmo2^CD2
SOX11^Lck;Lmo2^CD2

**Figure S5. Monitoring in vivo SOX11 activation and enhanced DN3 thymocyte development following combined SOX11 and LMO2 expression. (A)** Graph showing the percentage of GFP+ levels, used as a proxy for in vivo SOX11 activation, across different stages of T-cell differentiation in thymi of mice with T-cell restricted SOX11 overexpression *(SOX11*$^{Lck}$*)*. **(B)** Flow cytometric analysis of CD44 and CD25 expression within CD4$^{-}$CD8$^{-}$ DN thymocytes isolated from 8-week-old control, *SOX11*$^{Lck}$, *Lmo2*$^{CD2}$, or *SOX11*$^{Lck}$*;Lmo2*$^{CD2}$ mice. Cells were pregated for single, live CD90.2$^{+}$ cells. **(C)** Bar graphs quantifying the frequency of CD4$^{-}$CD8$^{-}$ DN thymocytes, expressed as a percentage of live cells, from mice expressing SOX11 alone *(SOX11*$^{Lck}$*)* or Lmo2 alone (*Lmo2*$^{CD2}$) or the combined transgenes (*SOX11*$^{Lck}$*; Lmo2*$^{CD2}$). Each dot represents a different mouse. A Mann-Whitney test was performed, and statistical significance is indicated as follows: * $P < 0.05$, ** $P < 0.01$, *** $P < 0.0001$.

**Figure S6.**

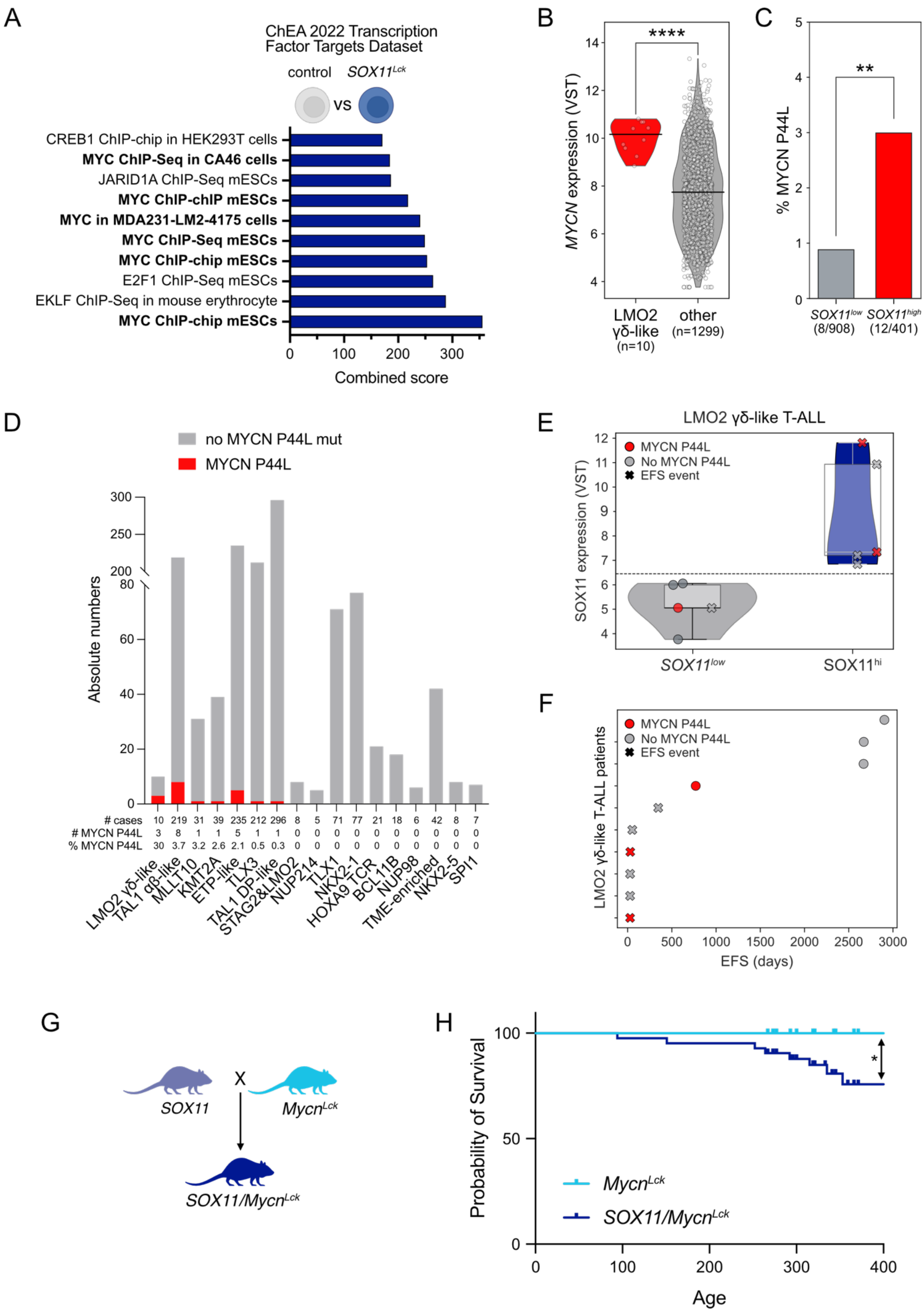
A
ChEA 2022 Transcription Factor Targets Dataset
control SOX11Lck
vs
CREB1 ChIP-chip in HEK293T cells
MYC ChIP-Seq in CA46 cells
JARID1A ChIP-Seq mESCs
MYC ChIP-chIP mESCs
MYC in MDA231-LM2-4175 cells
MYC ChIP-Seq mESCs
MYC ChIP-chip mESCs
E2F1 ChIP-Seq mESCs
EKLF ChIP-Seq in mouse erythrocyte
MYC ChIP-chip mESCs
Combined score
B
MYCN expression (VST)
****
LMO2 γδ-like (n=10)
other (n=1299)
C
% MYCN P44L
**
SOX11low (8/908)
SOX11high (12/401)
D
no MYCN P44L mut
MYCN P44L
Absolute numbers
### cases 10 219 31 39 235 212 296 8 5 71 77 21 18 6 42 8 7
### MYCN P44L 3 8 1 1 5 1 1 0 0 0 0 0 0 0 0 0 0
% MYCN P44L 30 3.7 3.2 2.6 2.1 0.5 0.3 0 0 0 0 0 0 0 0 0 0
LMO2 γδ-like, TAL1 αβ-like, MLLT10, KMT2A, ETP-like, TLX3, TAL1 DP-like, STAG2&LMO2, NUP214, TLX1, NKX2-1, HOXA9 TCR, BCL11B, NUP98, TME-enriched, NKX2-5, SPI1
E
LMO2 γδ-like T-ALL
MYCN P44L
No MYCN P44L
EFS event
SOX11 expression (VST)
SOX11low
SOX11hi
F
MYCN P44L
No MYCN P44L
EFS event
LMO2 γδ-like T-ALL patients
EFS (days)
G
SOX11
X
MycnLck
SOX11/MycnLck
H
Probability of Survival
*
MycnLck
SOX11/MycnLck
Age

**Figure S6. High SOX11 expression associates with MYCN P44L mutations and poor clinical outcome in LMO2 γδ-like T-ALL, while SOX11 and MYCN cooperate to drive T-ALL in vivo. (A)** Enrichment analysis bar graphs showing upregulated genes in *SOX11*$^{Lck}$ DN3 thymocytes compared to the WT condition. **(B)** Violin plot depicting *MYCN* expression after variance-stabilizing transformation (VST) in LMO2 γδ-like T-ALL (n=10) compared with all other T-ALL subtypes (n=1293). Significance was assessed using a two-sided Mann-Whitney U test ($P = 8.67 \times 10^{-5}$) **(C)** Bar graph showing the frequency of MYCN P44L mutations in *SOX11*$^{low}$ (8/908; 0,9%) and *SOX11*$^{high}$ (12/401; 3.0%) T-ALL cases from the Pölönen et al. cohort [1] . *SOX11*$^{high}$ was defined as VST expression > 4.5. Significance was assessed using Fisher's exact test (**, $P = 0.0065$). **(D)** Stacked bars show the absolute number of cases per T-ALL subtype[1] with MYCN P44L mutations (red) or without MYCN P44L mutations (grey), ranked by decreasing mutation frequency. The table below indicates the total number of cases, number of MYCN P44L-mutated cases, and percentage of MYCN P44L-mutated cases per subtype. **(E)** Violin plot of *SOX11* expression in LMO2 γδ-like T-ALL patients[1], stratified into SOX11$^{hi}$ and SOX11$^{low}$ groups using a within-subtype median split. Cases harboring the MYCN P44L-mutation are shown in red, non-mutated cases in grey, and patients who experienced an event are marked with crosses. **(F)** Swimmer plot of individual LMO2 γδ-like T-ALL patients[1] ranked by event-free survival (EFS) over time. Symbols and colors are as described in panel E. **(G)** Scheme depicting the breeding strategy to obtain *R26-Mycn*$^{tg/+}$*;LckCre*$^{tg/+}$ (*Mycn*$^{Lck}$) or *R26-Mycn*$^{tg/+}$*;R26-SOX11*$^{tg/+}$*;LckCre*$^{tg/+}$ (*SOX11/Mycn*$^{Lck}$) mice. **(H)** Kaplan Meier survival analysis of *Mycn*$^{Lck}$ and *SOX11/Mycn*$^{Lck}$ mice. Combined expression of SOX11 and MYCN significantly accelerates leukemia onset compared with MYCN overexpression alone (log-rank Mantel–Cox test; *, $P = 0.039$).

**References**

1. Polonen P, Di Giacomo D, Seffernick AE, et al. The genomic basis of childhood T-lineage acute lymphoblastic leukaemia. *Nature*. 2024;632(8027):1082-1091.
2. Inacio D, Amado T, Pamplona A, et al. Signature cytokine-associated transcriptome analysis of effector gammadelta T cells identifies subset-specific regulators of peripheral activation. *Nat Immunol*. 2025;26(3):497-510.

**Table S1 Genotyping primers**

| **Primer name** | **Primer sequence** | **PCR product** |
|---|---|---|
| *ROSA26 F* | 5'-CGAGCGGATAACAATTTCACA-3' | tg: 570 bp |
| *Ins R* | 5'-CCAAGCTTTTTCCCCGTATC-3' | |
| *ROSA26 5' F* | AAAGTCGCTCTGAGTTGTTAT | wt: 500 bp |
| *ROSA26 3' mut R* | GCGAAGAGTTTGTCCTCAACC | tg: 250 bp |
| *ROSA26 3' wt R* | GGAGCGGGAGAAATGGATATG | |
| *Tlox F* | 5'-ATCATGTCTGGATCCCCATC-3' | tg: 500 bp |
| *SOX11 R* | 5'-TCTTCCTGCCTTCGATCTTG-3' | |
| *iCre F (CD2-iCre)* | 5'-AGATGCCAGGACATCAGGAACCTG-3' | tg: 250 bp |
| *iCre R (CD2-iCre)* | 5'-ATCAGCCACACCAGACACAGAGATC-3' | |
| *Cre F (Lck-Cre)* | 5'-CGCCGCATAACCAGTGAAAC-3' | tg: 300 bp |
| *Cre R (Lck-Cre)* | 5'-ATGTCCAATTTACTGACCG-3' | |
| *CD2-Lmo2 tg F* | 5'-ATGTCCTCGGCCATCGAAAGGAAGAGCC-3' | tg: 480 bp |
| *CD2-Lmo2 tg R* | 5'-CCCATTGATCTTGGTCCACT-3' | |
| *CD2-Lmo2 wt F* | 5'-CTGTCCCCAACGGATTTCTA-3' | wt: 220 bp |
| *CD2-Lmo2 wt R* | 5'-ACGGTGAGGCCACTTGTATC-3' | |
| *Pten F* | 5'-TAGTGATAGAACGGAAGTCTTG-3' | fl: 1100 bp |
| *Pten R* | 5'-GATAAGTTCTAGCTGTGGTGG-3' | wt: 900 bp |
| *Mycn F* | 5'-ACCACAAGGCCCTCAGTACC-3' | tg: 300 bp |
| *Mycn R* | 5'-TGGGACGCACAGTGATGG-3' | |

**Table S2 qRT-PCR primers**

| Primer name | Primer sequence |
|---|---|
| *Floxed Stop F* | 5’-CACCTTCTACTCCTCCCCTA-3’ |
| *Floxed Stop R* | 5’-TACTTCCATTTGTCACGTCC-3’ |
| *R26-SOX11 F* | 5'- GTCAAGTGCGTGTTTCTGGA-3' |
| *R26-SOX11 R* | 5'-GACGTTGTAGCGTCTCAGCA-3' |
| *eGFP F* | 5’-CGACAACCACTACCTGAGCAC-3’ |
| *eGFP R* | 5’-CTTGTACAGCTCGTCCATGC-3’ |
| *Hprt1 F* | 5’-GGATTTGAATCACGTTTGTGT-3’ |
| *Hprt1 Rv* | 5’-TGGCAACATCAACAGGACTC-3’ |
| *Gapdh F* | 5’-CCCCAATGTGTCCGTCGTG-3’ |
| *Gapdh R* | 5’-GCCTGCTTCACCACCTTCT-3’ |
| *G6pdh F* | 5’-ATGCAGAACCACCTCCT-3’ |
| *G6pdh R* | 5’-TTCAACACTTTGACCTTCTCA-3’ |
| *Rpl13a F* | 5’-CCTGCTGCTCTCAAGGTTGTT-3’ |
| *Rpl13a R* | 5’-TGGTTGTCACTGCCTGGTACTT-3’ |
| *Hmbs F* | 5’-GAAACTCTGCTTCGCTGCATT-3’ |
| *Hmbs R* | 5’-TGCCCATCTTTCATCACTGTATG-3’ |
| *Tbp F* | 5’-TCTACCGTGAATCTTGGCTGTAAA-3’ |
| *Tbp R* | 5’-TTCTCATGATGACTGCAGCAAA-3’ |
| *Actb F* | 5’-GCTTCTAGGCGGACTGTTACTGA-3’ |
| *Actb R* | 5’-GCCATGCCAATGTTGTCTCTTAT-3’ |
| *Eef1a1 F* | 5’-TCGCCTTGGACGTTCTTTT-3’ |
| *Eef1a1 R* | 5’-GTGGACTTGCCGGAATCTAC-3’ |
| *Oaz1 F* | 5’-ATTGCTGTTTAAGATGGTCAGG-3’ |
| *Oaz1 R* | 5’-GGGGAGGTGACACTATTTTTCC-3’ |
| *Matr3 F* | 5'-TGGACCAAGAGGAAATCTGG-3' |
| *Matr3 R* | 5'-TGAACAACTCGGCTGGTTTC-3' |
| *B2m F* | 5'-CGGCCTGTATGCTATCCAGAA-3' |
| *B2m R* | 5'-GGCGGGTGGAACTGTGTTA-3' |
| *Ubc F* | 5'-AAAGCCCCTCAATCTCTGGAC-3' |
| *Ubc R* | 5'-TGCATCGTCTCTCTCACGGA-3' |

**Table S3 List of antibodies used for flow cytometry**

| Marker | clone | fluofore | dilution | company |
|---|---|---|---|---|
| ***TCRgd T-cell panel*** | | | | |
| Thy1.2/CD90.2 | 53-2.1 | BV500 | 1/250 | BD Biosciences |
| Live/dead stain | | PI | 1/100 | Merck Life Science |
| CD3 | 145-2C11 | PE | 1/200 | eBioscience |
| CD4 | RM4-5 | AF700 | 1/250 | eBioscience |
| CD8 | 53-6.7 | PE-Cy7 | 1/250 | eBioscience |
| CD25 | PC61 | BV421 | 1/250 | BD Biosciences |
| CD44 | IM7 | APC-Cy7 | 1/250 | BD Biosciences |
| TCRgd | GL3 | APC | 1/80 | Biolegend |
